# High Spatial Resolution Ambient Ionization Mass Spectrometry Imaging Using Microscopy Image Fusion Determines Tumor Margins

**DOI:** 10.1101/657494

**Authors:** Li-En Lin, Chih-Lin Chen, Ying-Chen Huang, Hsin-Hsiang Chung, Chiao-Wei Lin, Ko-Chien Chen, Yu-Ju Peng, Shih-Torng Ding, Ming-Yang Wang, Tang-Long Shen, Cheng-Chih Hsu

## Abstract

Mass spectrometry imaging (MSI) using ambient ionization technique enables a direct chemical investigation of biological samples with minimal sample pretreatment. However, detailed morphological information of the sample is often lost due to its limited spatial resolution. In this study, predictive high-resolution molecular imaging was produced by the fusion of ambient ionization MSI with optical microscopy of routine hematoxylin and eosin (H&E) staining produces. Specifically, desorption electrospray ionization (DESI) and nanospray desorption electrospray ionization (nanoDESI) mass spectrometry are employed to visualize lipid and protein species on mice tissue sections. The resulting molecular distributions obtained by ambient ionization MSI-microscopy fusion are verified with matrix-assisted laser desorption ionization time-of-flight (MALDI-TOF) MSI and immunohistochemistry (IHC) staining. Label-free molecular imaging with 5-μm spatial resolution can be acquired using DESI and nanoDESI, whereas the typical spatial resolution of ambient ionization MSI is ~100 μm. In this regard, sharpened molecular histology of tissue sections is achieved, providing complementary references to the pathology. Such a multimodality integration enables the discovery of potential tumor biomarkers. After image fusion, more than a dozen of potential biomarkers that could be used to determine the tumor margins on a metastatic mouse lung tissue section and Luminal B breast tumor tissue section are identified.

## 1. Introduction

Image fusion combining spatially-resolved data from multiple analytical tools has been utilized to generate high quality images for better human interpretation.^[1]^ A variety of image fusion methods, based on wavelet transform, artificial neural network, multivariate regression, and Pan-Sharpening methods, have been developed and successfully applied in different scientific fields.^[1–4]^ For instance, it has been largely used for remote sensing and object recognition by merging the satellite imaging with high resolution panchromatic image to harvest more information.^[1,5]^ One of the other successful applications of image fusion is in merging complementary medical images obtained with multiple modalities into one highly defined image for clinical analysis.^[4,6]^

Mass spectrometry imaging (MSI) provides spatially resolved chemical information on the surface of biological samples in a label-free manner, and has been profoundly used in preclinical, pharmaceutical and biological studies.^[7–16]^ Furthermore, ambient ionization mass spectrometry imaging methods are capable of providing spatially resolved chemical information of cells and tissues with minimal sample pretreatment under atmospheric condition. In specific, the biological tissue sections that subsequently used for conventional pathological and molecular staining can be directly implemented for ambient ionization MSI at their intact states. Desorption electrospray ionization (DESI) and nanospray-desorption electrospray ionization (nanoDESI), were two of the commonly used ambient ionization methods for *in-situ* analysis for different classes of compounds.^[10,11,17–23]^ MSI using DESI and nanoDESI is of great potential as a supplementary tool for pathological examinations in cancer diagnosis and has been largely used in determining the tumor margins.^[24–28]^ In previous reports, lipid species had been widely studied and were served as important biomarkers to identify the malignant and benign tumor tissue by DESI MSI.^[24,25]^ On the other hand, proteins were also reported to discriminate between normal tissues and tumors using nanoDESI MSI.^[11]^ However, one of the challenges for clinical study by DESI and nanoDESI MSI is its spatial resolving power.^[29–32]^

To visualize fine chemical details of the sample surface in its intact state comparable with conventional optical microscopy-based methods, the spatial resolution of ambient ionization MSI requires to improve.^[33]^ Several instrumental approaches to increase the spatial resolution of ambient ionization MSI have been reported. For example, a hybrid atomic force microscopy mass spectrometer was applied for mapping bacterial colonies at the submicrometer level under atmospheric pressure.^[34]^ On the other hand, laser desorption/ablation-based methods allow imaging at subcellular levels.^[35,36]^ DESI is currently the most widely recognized ambient ionization methods for MSI. Although its lateral resolution can reach to about 10 μm with optimal parameter settings,^[37]^ the typical resolution of DESI MSI is about 100 μm. NanoDESI is known for its ability of in situ protein MSI,^[10,11]^ but its spatial resolution is restricted by the instrumental design and occasional carried-over among pixels.

In addition to the instrumental approaches, numerical approaches, image fusion in particular, is an alternative strategy to surpass the inherent limitation of each ionization methods. This strategy allows us to achieve a higher spatial resolution without using custom instruments or specialized experimental setup. Recently, multivariate regression image fusion of optical and electron microscopy data with ultrahigh vacuum-based MSI, including matrix-assisted laser desorption ionization (MALDI) time-of-flight (TOF) and secondary ion mass spectrometry (SIMS) MSI, were reported to achieve sharpened molecular imaging with a spatial resolution at the cellular level on tissue sections.^[38–40]^ Meanwhile, these MSI have to be operated under high vacuum conditions and limit their compatibility with conventional imaging methods for pathological examinations.

In this study, an optical microscope was used to obtain images containing meticulous details of H&E stained mice tissue sections. Using multivariate regression algorithm, we fused ambient ionization MSI with an optical microscopy image so as to generate predictive MSI in different tissue types, and further used the fine-grained molecular mappings for cancer diagnosis. The workflow of our study is shown in **Figure 1** (see Supplementary Figure 1 for experimental setups), details of our results (see Supplementary Figure 2 for representative DESI and nanoDESI mass spectra), as well as experimental and data processing procedures (see Supplementary Figure 3 for the protocol of image fusion) are elaborated in the following sections.

**Figure 1.**
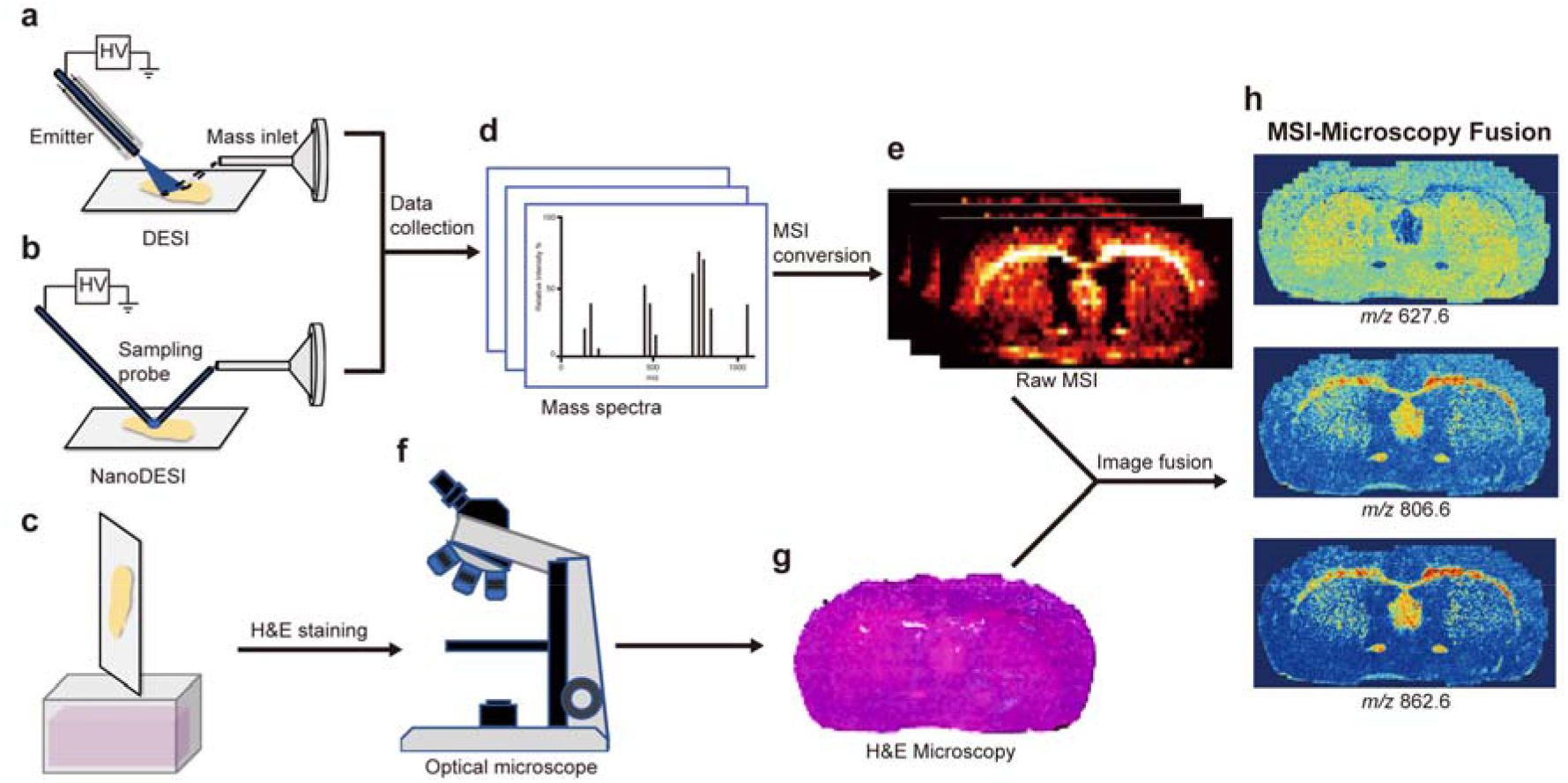
Illustration of the workflow to ambient ionization MSI-microscopy image fusion. (a) The scheme of DESI (desorption electrospray ionization) setup. (b) The scheme of homemade nanoDESI (nanospray desorption electrospray ionization) source. (c), (f), (g) H&E stained sections for optical microscopy image. (d) MS data collected from ambient ionization sources. (e) Alignment of DESI/nanoDESI MSI with optical microscopy images from (g). After MSI data collection from the ambient ionization sources, optical microscopies of H&E stained residue (for DESI) or adjacent (for nanoDESI) sections were employed for predictive chemo-spatial distribution based on statistic evaluations to generate high spatial resolution molecular imaging (h).

## 2. Results

### 2.1. Lipid Mapping by DESI MSI with Elevated Spatial Resolution

The results for the fusion of DESI MSI for mice brain and cerebellum coronal sections were shown in **Figure 2**, whereas the results for mice kidney sections were shown in **Figure 3** (see Supplementary Figure 4 and Supplementary Table 1 for compound identification using tandem mass analysis). For the raw lipid mappings in mice brain and cerebellum, as shown in Figure 2b and 2e, the distribution of two phospholipids species, phosphatidylethanolamine (PE P-18:0/20:5) and phosphatidylcholine (PC 18:0/20:1) at *m/z* 772.5 and 838.6, respectively, were revealed by our raw DESI MSI at 150-μm resolution. PE (P-18:0/20:5) were largely found in the grey matter, whereas PC (18:0/20:1) were measured in the white matter, and similar distributions of these two phospholipids species in mammalian brains had been reported using DESI MSI.^[41,42]^ Furthermore, in Figure 3b and 3f, PE (P-18:0/20:5) and PC (16:0/16:0) distributed in the cortex, while the PC (16:0/18:1) distributed in the medulla of the kidney, where these results were also similar to the previous studies.^[43,44]^ Although the DESI MSI provided molecular imaging at sub-tissue level, in which different lipid species showed a drastic difference in their distribution in the brain, morphological details were missing due to its limitation in spatial resolution. After the fusion of DESI MSI with the optical microscopic image of the H&E staining using the same tissue sections, predictive molecular imaging of the denoted phospholipids were generated, showing much improved morphological details compared with the raw DESI MSI. For example, the distribution of PE (P-18:0/20:5) from the raw DESI MSI only roughly revealed the hippocampus (Figure 2b), while the predictive image (Figure 2c) showed that PE (P-18:0/20:5) had lower abundance in the dentate gyrus. In the case of the cerebellum (Figure 2e and 2f), the originally pixelated molecular imaging was remarkably sharpened by image fusion, distinguishing between the cerebellar cortex and medulla. For the kidney sections in Figure 3c and 3g, clear boundaries between the outer stripe of the outer medulla (OSOM) and the inner stripe of the outer medulla (ISOM) were drastically revealed after the image fusion, and their distributions were in agreement with the results obtained by MALDI-TOF MSI using adjacent sections (Figure 3d and 3h).

**Figure 2.**
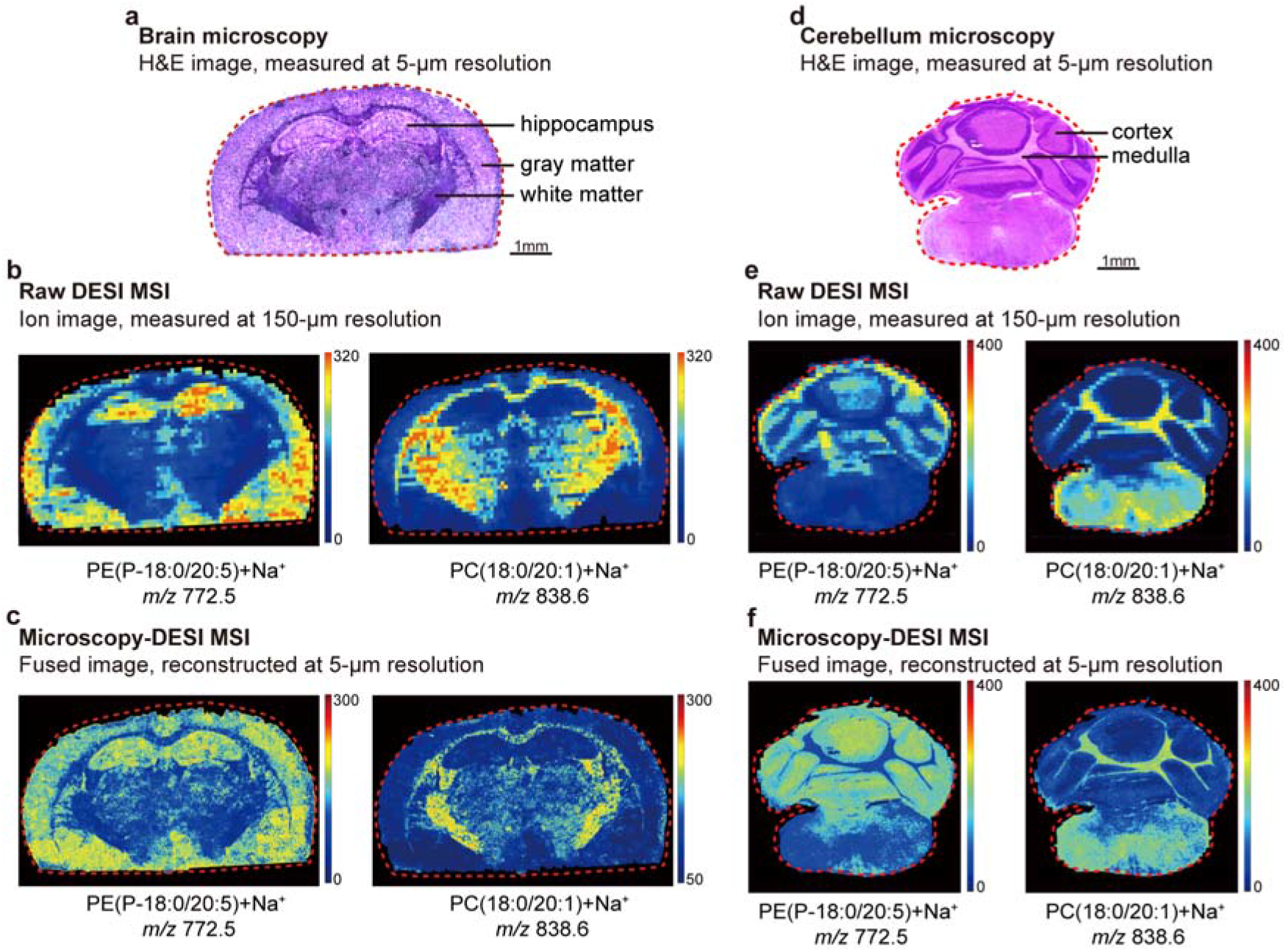
Molecular imaging of lipids in mice brain and cerebellum mapped by DESI MSI. (a) H&E stained mouse brain section. (b) Raw DESI MSI of lipid species. (c) Predicted high spatial resolution MSI of the mouse brain section after image fusion. (d) H&E stained mice cerebellum section. (e) Raw DESI MSI of lipid species. (f) Predicted high spatial resolution MSI of the mouse cerebellum section after image fusion.

**Figure 3.**
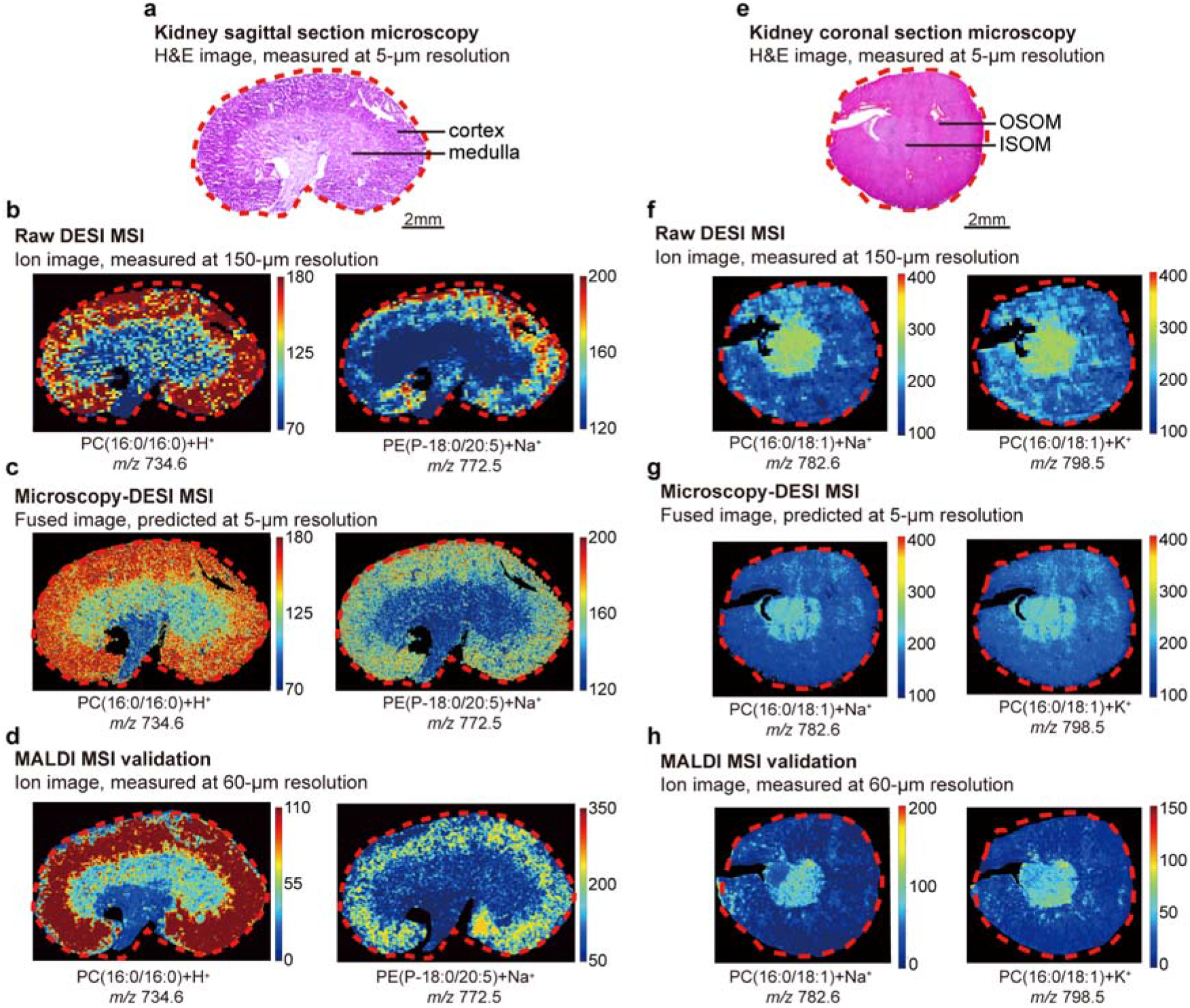
Molecular imaging of lipids in mice kidney sagittal and coronal sections by DESI and MALDI-TOF MSI. (a) H&E stained sagittal section of mouse kidney. (b) Raw DESI MSI of PC (16:0/16:0) and PE(P-18:0/20:5). (c) Predicted high spatial resolution MSI of lipids in the mouse kidney sagittal section after image fusion. (d) Lipid mappings of the adjacent kidney sagittal sections by MALDI-TOF MSI. (e) H&E stained coronal section of mouse kidney. (f) Raw DESI MSI of PC (16:0/18:1) ions. (g) Predicted high spatial resolution MSI in the mouse coronal section after image fusion. (h) Molecular imaging of the adjacent mouse kidney coronal section by MALDI-TOF MSI.

### 2.2. High-quality Protein Imaging by nanoDESI MSI

Unlike DESI, nanoDESI utilizes a micro-liquid junction sustained between two fused silica capillaries to desorb analyte compounds from the sample surfaces. This allows nanoDESI MSI to detect large biomolecules, such as proteins, with molecular weights up to 15 kDa.^[11,18,45,46]^ However, as the stability of the micro-junction is sensitive to the flatness of the tissue sections, the carry-over of the analyte compounds among neighboring pixels in nanoDESI MSI is thus more significant than in DESI MSI, causing artifacts to the images and making it challenging to be interpreted. To expand the utility of image fusion to protein imaging, we applied nanoDESI MSI to mice tissue sections. As shown in **Figure 4**, the spatial distribution of peak *m/z* 785.55 (+18 charge), annotated as myelin basic protein (MBP) using top-down tandem mass analysis (Supplementary Figure 5), was largely found on the mouse brain and mapped by nanoDESI MSI. Similar to the previous report, as MBP is highly expressed in the myelinated axons, it was profoundly observed in the white matter of the brain (Figure 4b).^[11]^ After image fusion with the H&E stained adjacent section, a high-resolution image of MBP was thus generated (Figure 4c). Remarkably, the severe tailing and cross-contamination of ion signals in the raw nanoDESI MSI due to carry-over effect was significantly eliminated in the predictive image. The distribution of MBP was well sharpened and in agreement with the histological features of the mouse brain section. In order to prove the genuineness of the prediction using image fusion, a high resolution MALDI-TOF MSI was also implemented to the second adjacent sections. The resulting MALDI-TOF MSI (Figure 4d) shows a very high similarity with the predictive image and thus verified that the pixel-wised carry-over was efficiently removed after image fusion.

**Figure 4.**
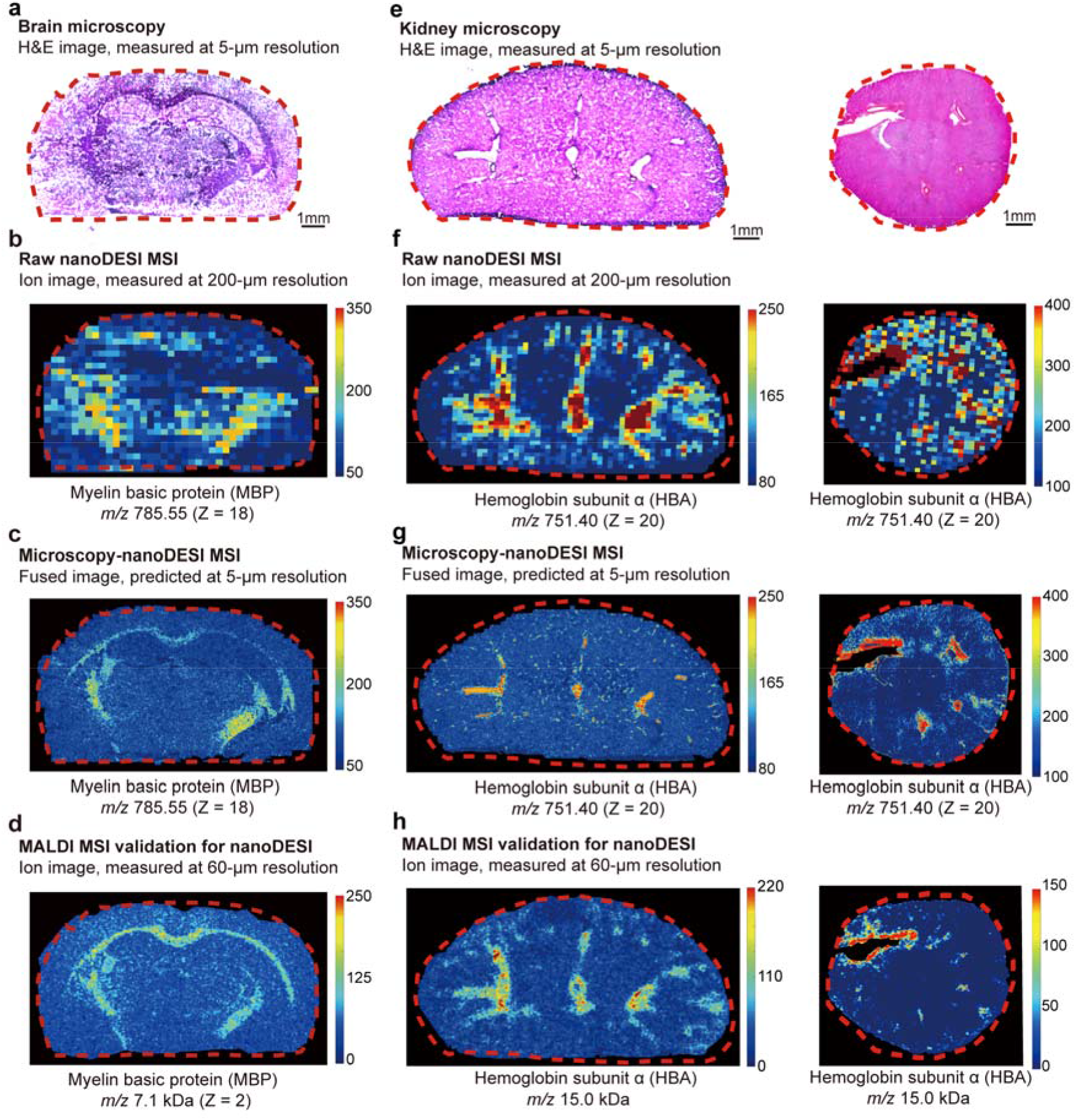
Molecular imaging of proteins in mice brain and kidney sections by nanoDESI MSI and microscopy image fusion. (a) H&E stained adjacent mouse brain tissue section of nanoDESI MSI. (b) Raw nanoDESI MSI of myelin basic protein (MBP) at ~200-μm spatial resolution. (c) Predicted high spatial resolution MBP distribution in the mouse brain after image fusion with microscopy image. (d) MALDI-TOF MSI of MBP in the adjacent mouse brain section to prove the efficacy of image fusion. (e) H&E stained adjacent mice kidney tissue sections of nanoDESI MSI. (f) Raw nanoDESI MSI of hemoglobin subunit alpha (HBA) at ~200-μm spatial resolution. (g) Predicted high spatial resolution HBA distribution in the mice kidney after image fusion with microscopy image. (h) MALDI-TOF MSI of HBA in the adjacent sections.

In addition, the spatial distribution of hemoglobin subunit alpha (*m/z* 751.40) was successfully mapped in sagittal and coronal kidney sections as shown in Figure 4f (see Supplementary Figure 6 for protein identification). The predictive image after fusion with H&E stained sections exhibits that a large amount of hemoglobin subunit alpha was still located in the blood vessels and not smeared during the cryo-sectioning (Figure 4g). The high-resolution MALDI-TOF MSI of the adjacent sections also verified this observation (Figure 4h). Notably, due to the fact that the histological features of kidney change after a few sections, a slight difference in the protein distribution obtained by nanoDESI MSI-microscopy fusion and MALDI-TOF MSI were expected.

### 2.3. Validations of Predicted MSI

We have demonstrated that DESI/nanoDESI MSI-microscopy image fusion provides an ability to resolve the molecular distribution of biomolecules on tissue sections at the cellular level (~5 μm), which is comparable with the traditional immunohistochemical approaches. However, immunohistochemistry (IHC) staining requires pre-requisite knowledge to the target compounds, proteins in specific, and does not allow label-free analysis. Thus, to further demonstrate the capability of DESI/nanoDESI MSI-microscopy image fusion in resolving the distribution of biomolecules in a label-free manner, a series of adjacent mice brain sections were prepared and sequentially analyzed with (1) DESI, (2) nanoDESI MSI-microscopy fusion, (3) MALDI-TOF MSI, and (4) IHC staining microscopy for lipid and protein imaging (**Figure 5**).

**Figure 5.**
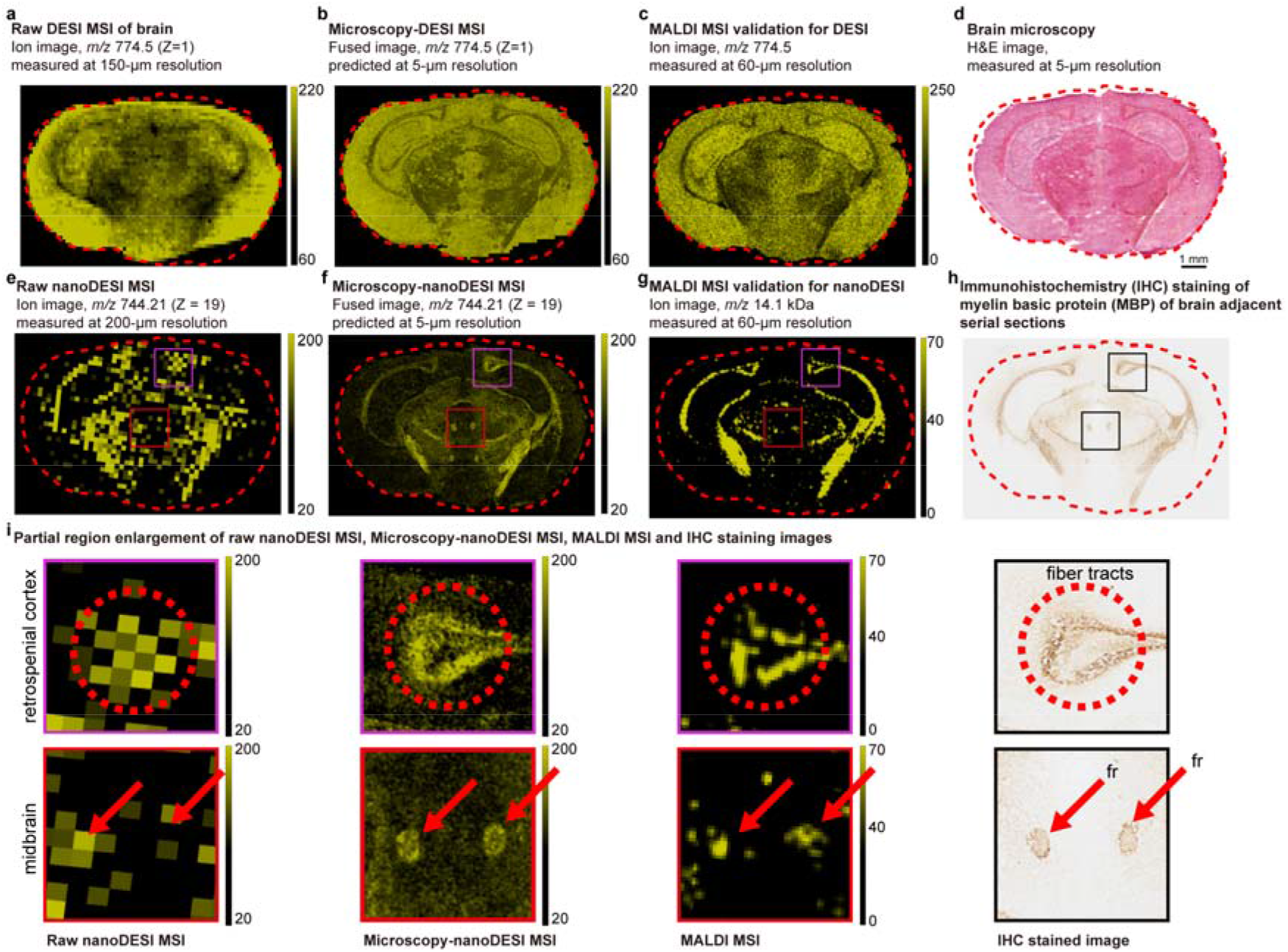
Verification of the efficacy of ambient ionization MSI-microscopy image fusion with MALDI-TOF MSI and immunohistochemistry (IHC) staining. (a) Raw DESI MSI of *m/z* 774.5 in the mouse brain section. (b) Predicted high spatial resolution MSI of *m/z* 774.5 after image fusion. (d) The distribution of *m/z* 774.5 obtained by MALDI-TOF MSI. (d) H&E stained serial section of the mouse brain. (e) Raw nanoDESI MSI of myelin basic protein (MBP) in the mouse brain section. (f) Predicted high spatial resolution MSI of MBP after image fusion with H&E stained microscopy image. (g) The distribution of MBP obtained by MALDI-TOF MSI. (h) IHC staining of MBP showing the same results by the predicted MSI. (i) Enlarged images of MBP distribution in the selected area of the mouse brain obtained by nanoDESI MSI, MSI-microscopy image fusion, MALDI MSI and IHC staining.

For lipid mapping shown in Figure 5a-c, the *m/z* 774.5 was observed in the region of the cerebral cortex by DESI imaging. However, the boundary was blurred and unclear by DESI MSI (Figure 5a) compared to the MALDI-TOF MSI (Figure 5c). Meanwhile, after image fusion was incorporated, the structure of the brain section was resolved in the predictive high spatial resolution molecular imaging (Figure 5b). For example, the hippocampus was unambiguously sketched in the mapping of this ion, showing comparable results with the MALDI-TOF MSI.

For protein detection shown in Figure 5e-i, MBP (*m/z* 744.21, +19 charge) was visualized by nanoDESI MSI. The nominal spatial resolution (determined by the speed of translational stage and rate of data collection) of raw nanoDESI MSI was 200 μm, and the tissue histological details were relatively difficult to be recognized due to the limited spatial resolution as a result of the large micro-junction radius and carry-over. In the raw nanoDESI MSI (Figure 5e), the histochemical relationship of MBP was ambiguous, especially at the brain stem area. In Figure 5i, the images at the retrosplenial cortex (purple) and the midbrain (red) are enlarged. After image fusion, MBP was predicted to be localized at the fiber tracts and fasciculus retroflexus (fr). Remarkably, similar results were obtained with MALDI-TOF MSI (Supplementary Figure 8). MBP is one of a major component of the myelin sheath. As a result, MBP can be easily detected in the white matter of a brain. To verify the result resolved by our nanoDESI MSI, high-resolution microscopic image of MBP was obtained by IHC staining of the neighboring section.^[47]^ Unambiguously, we were able to observe highly comparable distributions of MBP from the results of nanoDESI MSI-microscopy image fusion and the conventional IHC staining, while eliminating complex and time-consuming sample pretreatment and the need for specific antibodies.

### 2.4. Image Fusion of DESI MSI for Improved Cancer Diagnosis

DESI MSI had been widely applied to study various cancer models,^[23–26,28]^ allowing us to delineate the tumor margins on a heterogeneous section at the chemical level. Although ambient ionization MSI provides a simple and rapid molecular evaluation of clinical specimens,^[48]^ its limited spatial resolution makes it difficult to assess small-size tumors, especially at the early stage or metastatic cancer. Failure in distinguishing cancer cells from benign tissue increase the likelihood for a second operation.^[49]^ Thus, we seek to implement DESI MSI-microscopy image fusion to obtain greater details of molecular distribution in the cancer section. To evaluate its capability of predicting the precise cancerous margin, we applied image fusion to obtain fine spatial resolution molecular imaging in breast cancer metastatic lung tissue sections (**Figure 6**). The raw and predicted distributions of several phospholipids, including phosphatidylethanolamine (PE) and phosphatidylinositol (PI) were compared with the pathological examination (see Supplementary Figure 8 and Supplementary Table 2 for compound annotations). As elaborated in Figure 6d, many of the tiny tumors are about the similar sizes with imaging pixels in the raw data, and thus no enough statistical estimations could be made to judge whether the denoted lipid species were observed or not. Meanwhile, the marker lipid, e.g. PI (20:4/18:0), was predicted to appear in the selected tiny tumors, suggesting that imaging fusion could be utilized to improve the performance of the ambient ionization MSI-based cancer diagnosis.

**Figure 6.**
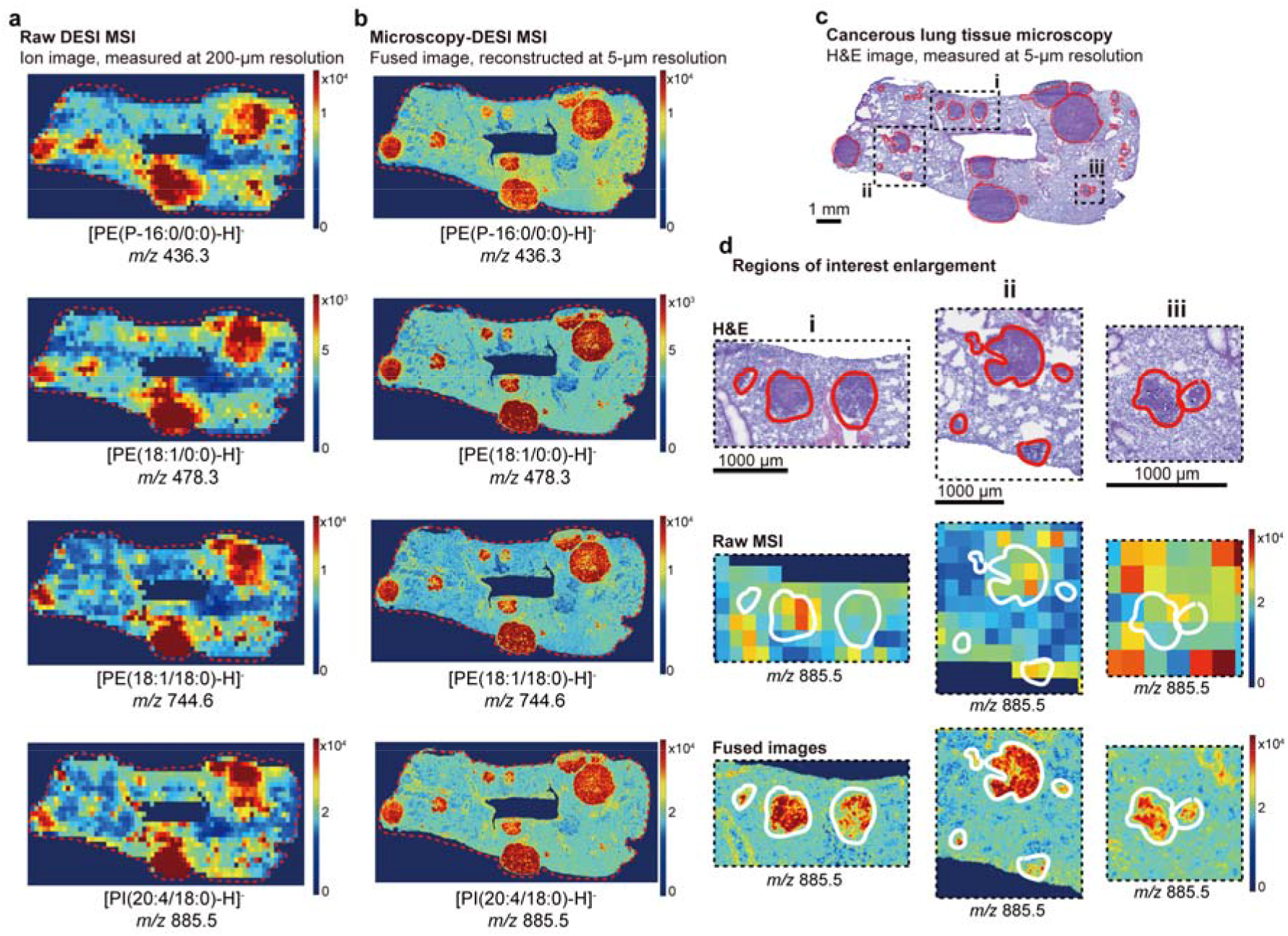
Ambient ionization MSI-microscopy image fusion reveals the biomarker lipids to the metastatic tumors in a mouse lung tissue. (a) Raw DESI-MSI of the representative lipid species. (b) Predicted high spatial resolution mappings of the biomarker lipid after microscopy-MSI image fusion. (c) H&E stained lung tissue section with the cancerous regions circled in red. (d) Enlarged images of microscopy image, raw DESI MSI and MSI-microscopy image of [PI(20:4/18:0)-H]^−^ species in the selected regions of the metastatic mouse lung section. In addition, in order to determine whether the molecular distributions correlate with the cancer region in the tissue section, and to further verify if the predicted DESI MSI has greater performance in the mining of biomarker compounds, the receiver operating characteristic (ROC) curves were plotted for each raw molecular imaging (Supplementary Figure 9). ROC curve is a binary classifier effective in evaluating the diagnostic ability of tests across all the possible threshold values.^[50]^ It has been profoundly used in evaluating tests for detections of cancers over the past decades.^[51–53]^ In our study, spatial-chemical results of the metastatic lung section obtained by DESI MSI were compared with the pathological evaluations. We would like to highlight here that since DESI MSI is tissue-friendly, the same section could be reused for H&E staining and sent out for the routine pathology assessment. Subsequently, the ROC curves for each predicted molecular imaging were calculated and plotted as in Supplementary Figure 9, in which the ion with a larger area under curve (AUC) of the ROC curve indicates that the ion has greater potential to serve as an effective biomarker.

To demonstrate the capability of this technique in analyzing human clinical samples, we further applied image fusion in a DESI MSI of a Luminal B human breast cancer tissue section (**Figure 7**) and plotted the ROC curves (Supplementary Figure 10). After image fusion, the AUC of 6 ion species increased significantly from ~0.69 to ~0.80 (see Figure 7). The predicted molecular images indicated that the signal intensities of these ions were higher on the cancerous region, and thus could potentially serve as biomarkers for cancer diagnosis.

**Figure 7.**
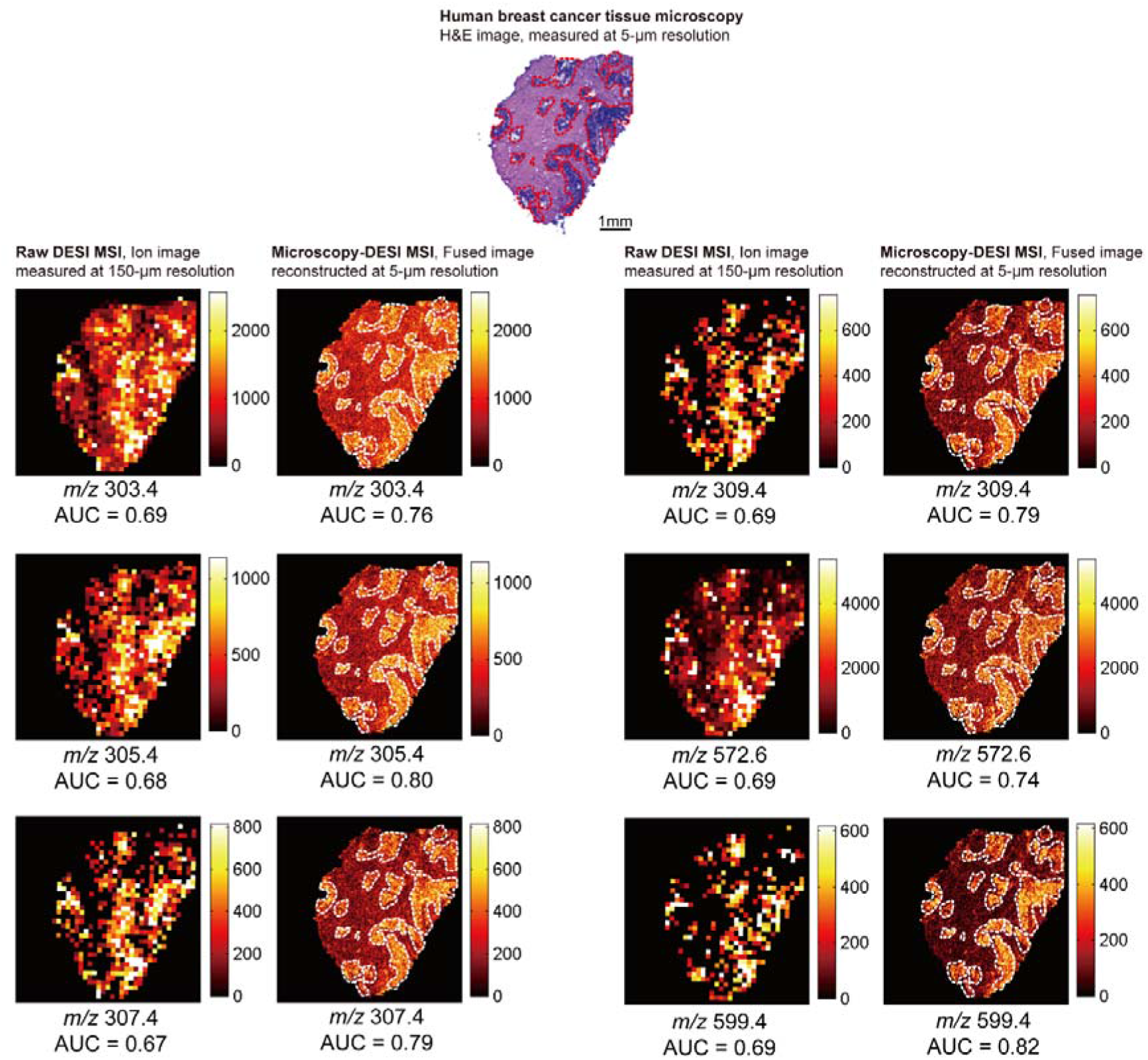
Ambient ionization MSI-microscopy image fusion reveals the potential biomarker lipids of the Luminal B human breast cancer tissue to determine the tumor margins. The cancerous regions are circled by red (in the H&E image) and white (in the fused image) dashed lines.

On the other hand, ions having AUC of the ROC curves around 0.5, such as *m/z* 780.6 in **Figure 8**a, represents that the distributions of the ion species were irrelevant to the presence of the tumors. In this regard, we defined the ions with AUC values > 0.7 as the potential biomarkers that can represent the cancerous region in the tissue sections. Using this criterion, in the metastatic lung tissue section, we found 16 ions with AUC of the ROC curves > 0.7, and the *m/z* values of these potential biomarkers were listed in Supplementary Table 3. For instance, as shown in Figure 8a, the raw DESI MSI of *m/z* 743.4 ion had an AUC value ~ 0.77, indicating a high spatial correlation between the ion signal and pathological interpretation. However, in the raw DESI MSI, the distribution of the ions had obscure outlines and thus it is difficult to differentiate the malignant tissue from the normal one. Therefore, some potential biomarkers were neglected. As have been demonstrated previously, the outline of the ions was sharpened after image fusion, making it easier to determine the tumor margins. In specific, as shown in Supplementary Figure 9 and 10, the ROC curves were bent more off-diagonally, meaning that the strengths of the classifiers were increased. More importantly, many of the ineffective markers (ions with AUC values between 0.5 to 0.7) in raw MSI were then be able to be recognized as effective biomarkers (AUC values > 0.7) in the predictive MSI. For example, as shown in Figure 8a, the AUC of *m/z* 724.6 ion increased from 0.65 to 0.79 after image fusion, making it a new potential biomarker of the tumor. This demonstrated the power of microscopy-DESI MSI in consolidating the effectiveness of molecular classifier to cancer assessment. In fact, by utilizing the greatly improved molecular mapping of high resolution microscopy-DESI MSI, 11 new ion species (equivalent to ~68% of the original number) were recognized as potential biomarkers for the metastatic cancer, and 6 new ion species (equivalent to ~22% of the original number) for the Luminal B breast cancer. The list of these biomarkers was elaborated in the supporting information Supplementary Table 3 and Supplementary Table 4.

**Figure 8.**
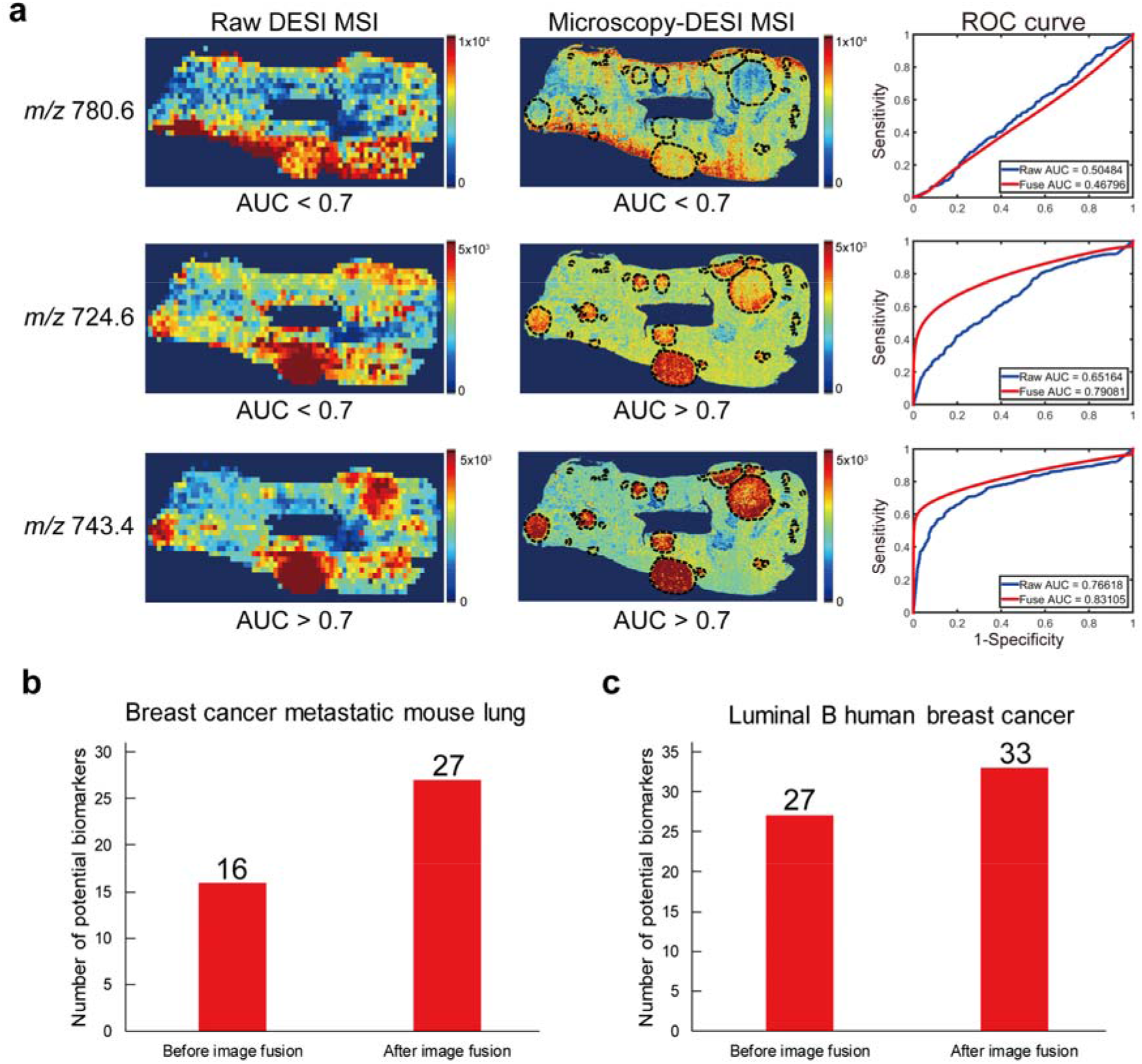
MSI-microscopy image fusion assists in searching potential biomarkers for tumor margin determination. (a) Raw and predicted MSI of the representative ions for breast cancer metastatic mouse lung tissue, with their corresponding ROC curves and AUC values presented in the right. The cancerous regions were circled in black. Ions that were ineffective in classifying tissue types, e.g. *m/z* 780.6, their AUC of ROC curve remained at ~0.5 before and after image fusion. However, for ions with higher AUC that served as classifiers, e.g. *m/z* 724.6 and 743.4, significant increases of AUC were obtained from the sharpened MSI after image fusion. Notably, some of the ions (such as *m/z* 724.6) with AUC below 0.7 in the raw MSI, had AUC exceeding 0.7 after image fusion and were subsequently considered as potential biomarkers. (b) The total numbers of potential biomarkers (AUC > 0.7) of the breast cancer metastatic mouse lung tissue increased from 16 in raw MSI to 27 in the predicted MSI (equivalent to 68% increased). (c) The total numbers of potential biomarkers (AUC > 0.7) of the Luminal B human breast cancer tissue increased from 27 in raw MSI to 33 in the predicted MSI (equivalent to 22% increased). *Note*: The ROC curves and corresponding AUCs of all ion images obtained in the metastatic mouse lung section were elaborated in the Supplementary Figure 9; and the AUCs of ions obtained in the human breast cancer tissue were shown in the Supplementary Figure 10.

## 3. Discussion

The development of ambient ionization MSI in the past decade has pushed the boundary of cancer diagnosis to the chemical level.^[10,15,23–26,28,54]^ As the tissue samples can be investigated directly under atmospheric pressure with only minimal sample preparation, it allows the integration of pathological examination and biomolecular mapping in a single section. However, due to its instrument limitation in spatial resolving power, much of the biological information is missing in ambient ionization MSI. Although the near-cellular resolution could be achieved with optimal instrument settings, to obtain the similar ion intensity and signal-to-noise ratio as in low resolution MSI, the acquisition time required for MSI becomes enormous because fewer target compounds are left in smaller pixels. In addition, when the spatial resolution of an MSI is doubled, the overall scanning time is consequently quadrupled. Longer scanning time also represents more serious compound degradation during imaging. Since the chemical compositions of the sample are altered as it is exposed to the ambient environment, minimizing the acquisition time of MSI is thus crucial.^[55]^ Utilizing numerical image fusion allows not only an improved MSI quality but also a higher data throughput, since intensive scanning is no longer required. As such, an optical microscopy-ambient ionization MSI was proposed and implemented to generate predictive MSI with a lateral resolution of the conventional optical microscopic data. We were able to extract the biological details to make a chemically-rich, high spatially-resolved image from a single tissue section, that is compatible with both the routine histological protocol and MSI.

In this paper, we reported the fusion of MSI and H&E images with multivariate regression. As the machine learning strategies become more affective in generating predictive images, we believe that applying these strategies (such as cycle GANs) in translating the MSI data into detailed molecular imaging is worthwhile to explore.^[56]^

## 4. Conclusion

In this study, we showed that the image fusion of DESI and nanoDESI MSI with H&E staining microscopy image was able to generate predictive imaging of biomolecules in various tissue sections. The morphological details were revealed in the high resolution MSI on mice brain, cerebellum, and kidney sections. More importantly, the predictive lipid and protein images were further validated by the comparison with MALDI-TOF MSI and conventional IHC staining microscopy. The unambiguous prediction by microscopy-ambient ionization MSI allowed a more precise determination of tumor margins and deeper mining of cancer biomarkers. Using a metastatic lung tissue section and a human breast cancer tissue section as the proof-of-concept, we disclosed more than 10 new ion species that could serve as potential tumor biomarkers. As it is nicely compatible with the routine histological protocols and pathological evaluations, image fusion of ambient ionization MSI with H&E staining microscopy images showed a great possibility for future applications in clinical diagnosis. We believe that our results have contributed towards this end.

## 5. Experimental Section

### Tissue sample preparation

Mice brain and kidney tissues were purchased from BioLASCO Taiwan Co., Ltd. Gender and age of the ICR mice were not specified. Metastatic lung tissues derived from MMTV-PyMT breast cancer mice model were collected from the laboratory of Dr. Tang-Long Shen (National Taiwan University) and the experimental details were described elsewhere.^[57,58]^ All experimental procedures were handled in accordance with the protocols and the ethical regulations approved by the Institutional Animal Care and Use Committee of National Taiwan University (IACUC approval NO. NTU104-EL-00003). The intact organs were harvested as soon as euthanasia. The Luminal B human breast cancer tissue were collected from National Taiwan University Hospital. All procedures on human tissue were performed with the approval of the Research Ethics Committee B of National Taiwan University Hospital (NTUH-REC approval No. 201812125RINB). The intact organs and tissue samples were stored under −80°C prior cryo-sectioning. For tissue cryo-sectioning, organs and tissue samples were flash-frozen using liquid nitrogen and kept under −20°C in the cold tome (LEICA, CM1900) in order to reach the optimal temperature before sectioning. The samples were sectioned to 14-μm thick sections and thaw-mounted onto the slides and stored at −80°C prior to analysis without fixation. Slides used for nanoDESI MSI analysis were regular plain glass without any coating. For DESI, H&E staining and immunostaining, tissues were thaw-mounted onto silane coated slides; while the slides for MALDI-TOF analysis were coated with indium tin oxide. Before nanoDESI MS interrogation, tissue sections were dried in a desiccator for ~1 hour and then rinsed with 50 ml of chloroform for 1 min. Adjoining tissue section pretreatment for MALDI-TOF validation is as described in the following. The matrix application was achieved by the sublimation method, which has been described elsewhere,^[59]^ followed by a recrystallization step.^[60]^ The sublimation apparatus was purchased from Singlong (Taichung, Taiwan) and placed in a sand bath on a hot plate while applying matrix. The brain sections were adhered to ITO-coated glass slides by a conductive tape and stored under −80°C before applying matrix. Sublimation was performed using 2,5-dihydroxyacetophenone (2,5-DHA). The matrix sublimation and application were performed at 110 °C with a 0.7 Torr vacuum for 10 minutes. Amount of the applied matrices were determined by the exposure time. The matrix-coated samples were rehydrated with 50% TFA solution in an incubator at 37°C for 4 minutes. A sonication step was added to increase the signal for protein analysis.^[61]^ The sonication was operated by Elmasonic S 30 H ultrasonicator under continuous mode with a frequency of 37 KHz. As for DESI MS interrogation, tissue sections were dried in a desiccator for ~1 hour, no further sample pretreatment was conducted before analysis.

### Ambient ionization mass spectrometry imaging

For DESI MSI, commercial DESI source (Prosolia Inc., IN, USA) was mounted to Orbitrap Elite to conduct the MSI measurements (mass range 200 to 1000 *m/z*). The gas pressure of nitrogen was set at 150 psi, the angle of the spray head was set to 55°, the flow rate of the solvent (DMF:ACN = 1:1) was 2 μl/min and the voltage was 3.5 kV. The nanoDESI system was modified from the commercial DESI platform (mass range 700 to 800 *m/z*), in which two flame-pulled fused capillaries (O.D: 360μm, I.D: 250 μm) were implanted and used to substitute the original DESI emitter. The solvent delivery (primary) capillary (65% ACN with 1% formic acid), was applied with a high voltage (2.5kV) to extract and ionize the compound on the surface of the sample. The secondary capillary was used to deliver the extracts from the sample to the mass spectrometer. The MSI experiments were conducted using Prosolia’s DESI 2D system. The scanning rates of the motor stage for DESI and nanoDESI MSI were set at approximately 150 μm s^−1^ and 30 μm s^−1^, respectively. The acquired raw data were then imported into Firefly™ 2.2 for data conversion, then we imported the converted data into BioMAP to obtain the final DESI and nanoDESI MSI. For H&E staining, the tissue slides were rinsed with 70% EtOH and then 100% EtOH for 30 sec each and allow dry under vacuum. The hematoxylin staining was applied under 60°C for 40 sec. After hematoxylin staining, the slides were rinsed with H_2_O, then rinsed with acidified EtOH 0.3% for 3 sec, then rinsed with H_2_O. The bluing up was achieved by 1% NH4OH and finally H_2_O under room temperature. Then the slides were dipped into the eosin stain for 20 sec, then the slides were rinsed with H_2_O, then 80% EtOH, then 90% EtOH, and finally 99% EtOH at room temperature.

### Data preprocessing for imaging fusion

The MSI data was converted into a *.img file by Firefly™ 2.2 and exported into a built-in Matlab package MSiReader. Each dataset was overlaid with the H&E image and the MSI data in MSiReader to determine the region of interests.^[62]^ The mass spectral signals were binned into 2,000 features (equivalent to 0.4 *m/z* each for DESI MSI and 0.05 *m/z* each for nanoDESI MSI) and summed before exported as a uniformly spaced text file. The microscope images were collected by an optical microscope (WHITED INC., Taipei, Taiwan) with a PSC600-05C digital camera (OPLENIC CORP., USA) and the image data was processed by AOR AJ-VERT. The microscopy images were then exported as a uniform data array by the home-built Matlab script. An affine transform matrix that can describe the spatial relationship between MSI data and microscopy data was calculated using a second home-built Matlab script. Subsequently, the best fitting between the MSI data and the microscopic data could be found to ensure the optimal alignment. The processed data was exported by in-house generated Matlab script and imported into “Molecular image fusion system” under the command-line interface to generate the predictive MSI datasets.^[38]^ The output high resolution MSI with reconstruction score >75% were exported for further AUC ROC curves analysis.

### Receiver operating characteristic curve

The ROC curves were constructed by our home-built MATLAB code. The ROC curve is given by the corresponding values of the sensitivity and the (1-specificity) at various ion intensity thresholds for individual *m/z* to determine the binary classification of tissue. If the intensity of a specific *m/z* peak in a particular pixel was higher than the intensity threshold, the pixel was then considered as “Cancer”. Otherwise, the pixel was assigned as “Normal”. The resulting sets of assignments were compared, pixel by pixel, with the labels of the H&E stained metastatic lung tissue image evaluated by the pathologist to calculate the sensitivity and the specificity of each ion. The sensitivity (true positive rate) was defined as the ratio of “the number of Cancer pixels determined by both MSI and the pathologist” to “the number of Cancer pixels labeled by the pathologist”; the (1-specificity), or false positive rate, was defined as the ratio of “the number of Cancer pixels determined by MSI in the normal tissue region” to “the number Normal pixels labeled by the pathologist”. The ROC curves of an *m/z* peak were sketched by setting the intensity threshold from 0% to 100% of the highest intensity of the ion in all pixels.

### Immunofluorescence staining

All the procedures for immunostaining were based on the protocols of the commercial kit (TAHC03, BioTnA, Kaohsiung, Taiwan). After immunostaining, the slides were mounted and digitized with a Motic Easyscan Digital Slide Scanner (Motic Hong Kong Limited, Hong Kong, China) at ×40 (0.26 μm/pixel) with high precision (High precision autofocus). Motic Easyscan whole-slide images were viewed with DSAssistant and EasyScanner software at Litzung Biotechnology INC (Kaohsiung, Taiwan).

### Matrix-assisted laser desorption/ionization time-of-flight mass spectrometry imaging

MALDI-TOF MSI results were acquired using MALDI-TOF/TOF mass spectrometry (Autoflex Speed MALDI TOF/TOF system, Bruker Daltonics). The instrument was equipped with the third harmonic of Nd:YAG SmartBeamTM-II laser (355nm). Imaging spectra were recorded and processed by FlexControl 3.4 and FlexImaging 3.0 (Bruker Daltonics). The spectra were acquired in positive polarity with pixel-to-pixel resolution of 60 μm using the following parameters: laser attenuator offset at 80% of the maximum power; laser operating power at 90% under linear mode for protein analysis and 80% under reflectron mode for lipid analysis with smartbeam parameter at 2_small; laser repetition rate at 1 kHz; acquisition shots accumulated to 1,000 shots per pixel for imaging analysis. The resulting imaging spectra were processed using TopHat baseline subtraction and normalized to the total ion counts per pixel.

### Mass spectrometry-based molecular identifications

Protein ions generated by nanoDESI source were directly introduced to the LTQ Orbitrap Elite for top-down tandem mass analysis^[45,46]^. Protein ions of interest were chosen as the mass center of 5-*m/z* isolation window with an activation energy Q of 0.25 and utilize collision energy of 30%. The data was imported into Prosight PTM for identification^[63,64]^. Lipid species in the brain tissue was extracted with MTBE methods for further tandem mass analysis^[65]^. The HPLC-MS/MS analysis was performed using HPLC (LC-20AD, Shimadzu, Tokyo, Japan), coupled with Orbitrap Elite (Thermo Scientific). HPLC experiments were performed using C18 column (100*2.1 mm, 3.5 μm, Agilent) and following the gradient elution: mobile phase A = water with 0.1% formic acid (v/v); mobile phase B = acetonitrile and isopropanol (10:90, v/v) with 0.1 % formic acid (v/v); elution profile = 0.0-5.0 min (40% mobile phase B); 5.0-35.0 min (40-90% mobile phase B); 35.0-50.0 min (90% mobile phase B), column oven at 25°C, volume injection 10 μL and flow rate of 0.15 mL/min. Mass spectrometry acquisition parameters were as followed: positive ions mode, heater temperature 180 °C, sheath gas flow rate 35 arb, auxiliary gas flow rate 10 arb, sweep gas flow rate 10 arb, spray voltage 3.5 kV and capillary temperature 350°C. CID fragmentation was performed for targeting ion peaks observed in DESI analysis with collision energy of 30% and an activation Q of 0.25. The mass spectra were collected in FT mode with 30,000 resolving power. The mass spectral analysis was processed by Xcalibur QualBrowser.

## Supporting Information

Supporting Information is available online.

## Acknowledgements

This research was supported by Ministry of Science and Technology (MOST), R.O.C. (Grant nos.: MOST 106-2113-M-002-013-MY2, 107-2321-B-001-038-, and 108-2636-M-002-008-), and Center for Emerging Materials and Advanced Devices, National Taiwan University (NTU) (Grant nos.: NTU-ERP-108L880116). Li-En Lin was supported by MOST grant 106-2813-C-002-136-M. The instrument support from NTU Mass Spectrometry Platform was acknowledged. The laboratory services of histology and pathology by the experienced veterinary pathologist, Hao-Kai Chang, at Litzung Biotechnology Inc., Taiwan, was acknowledged.

## The table of contents entry

Fusion of ambient mass spectrometry imaging with microscopic image allowed label-free imaging of molecular distribution at 5-μm spatial resolution. Detailed images of protein and lipid species are mapped and confirmed. With the minimum sample preparation, capability of providing detailed distribution, and increased diagnostic ability, this techniques is applicable to cancer diagnosis.

## ToC figure

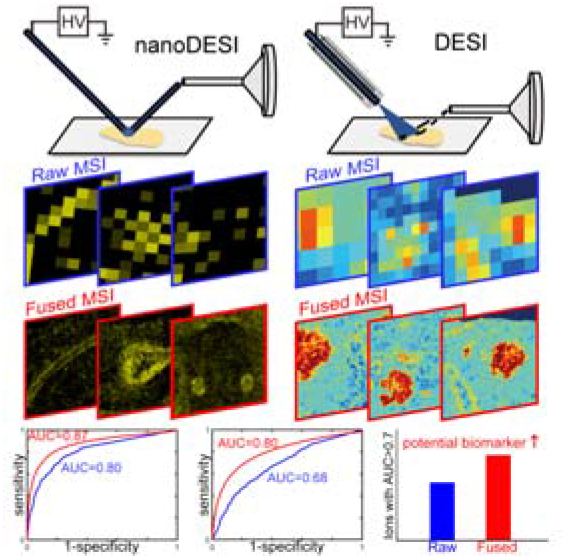

