## supplementary information for "High Spatial Resolution Ambient Ionization Mass Spectrometry Imaging Using Microscopy Image Fusion Determines Tumor Margins"

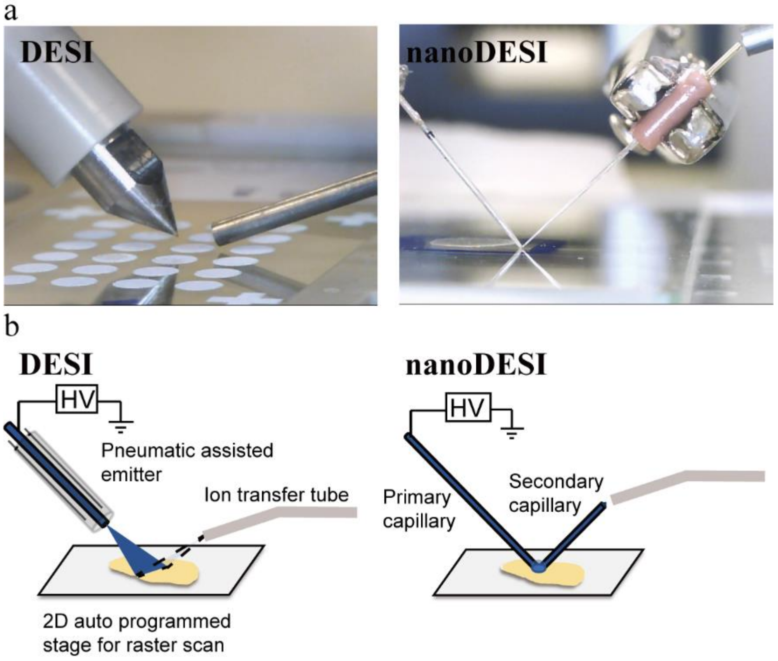


**Supplementary Figure 1.** Experimental setups of ambient ionization sources and 2-D translation mass spectrometry imaging (MSI) platform. (a) Photographs of the experimental setup. (B) Schematic illustrations of the setup.

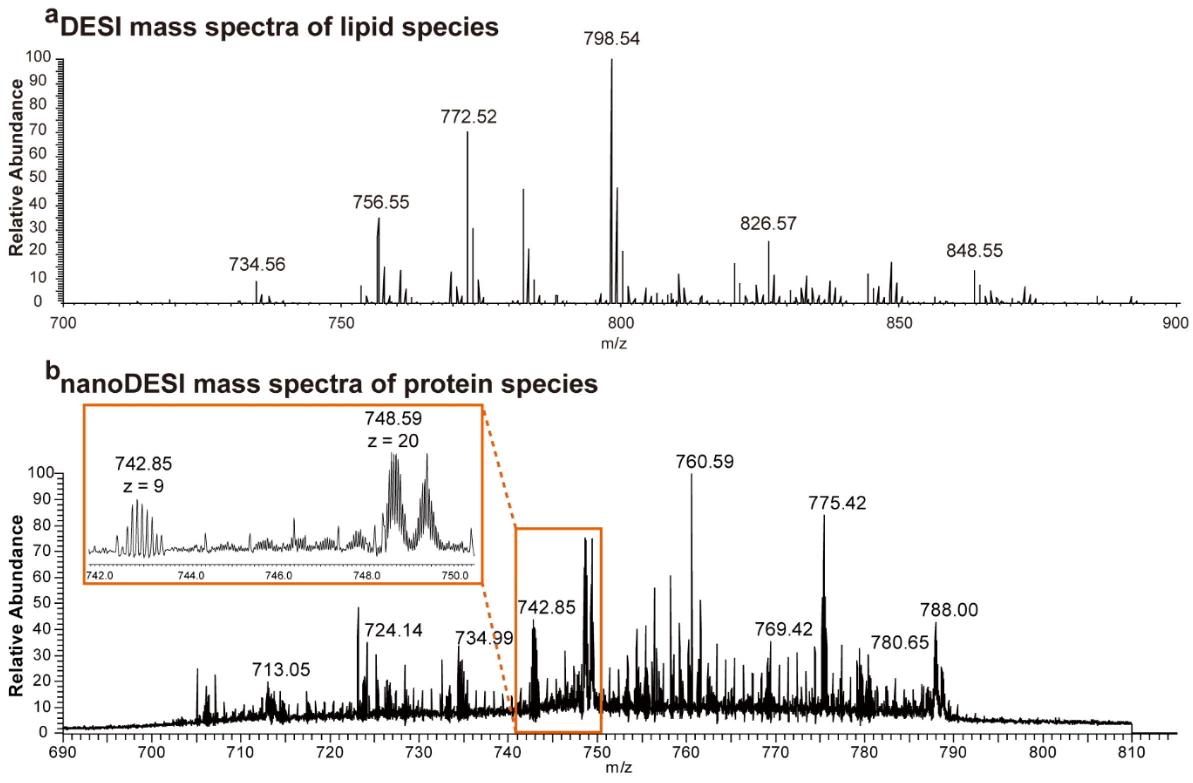


**Supplementary Figure 2.** Representative mass spectra obtain by DESI and nanoDESI MSI on mice brain sections.

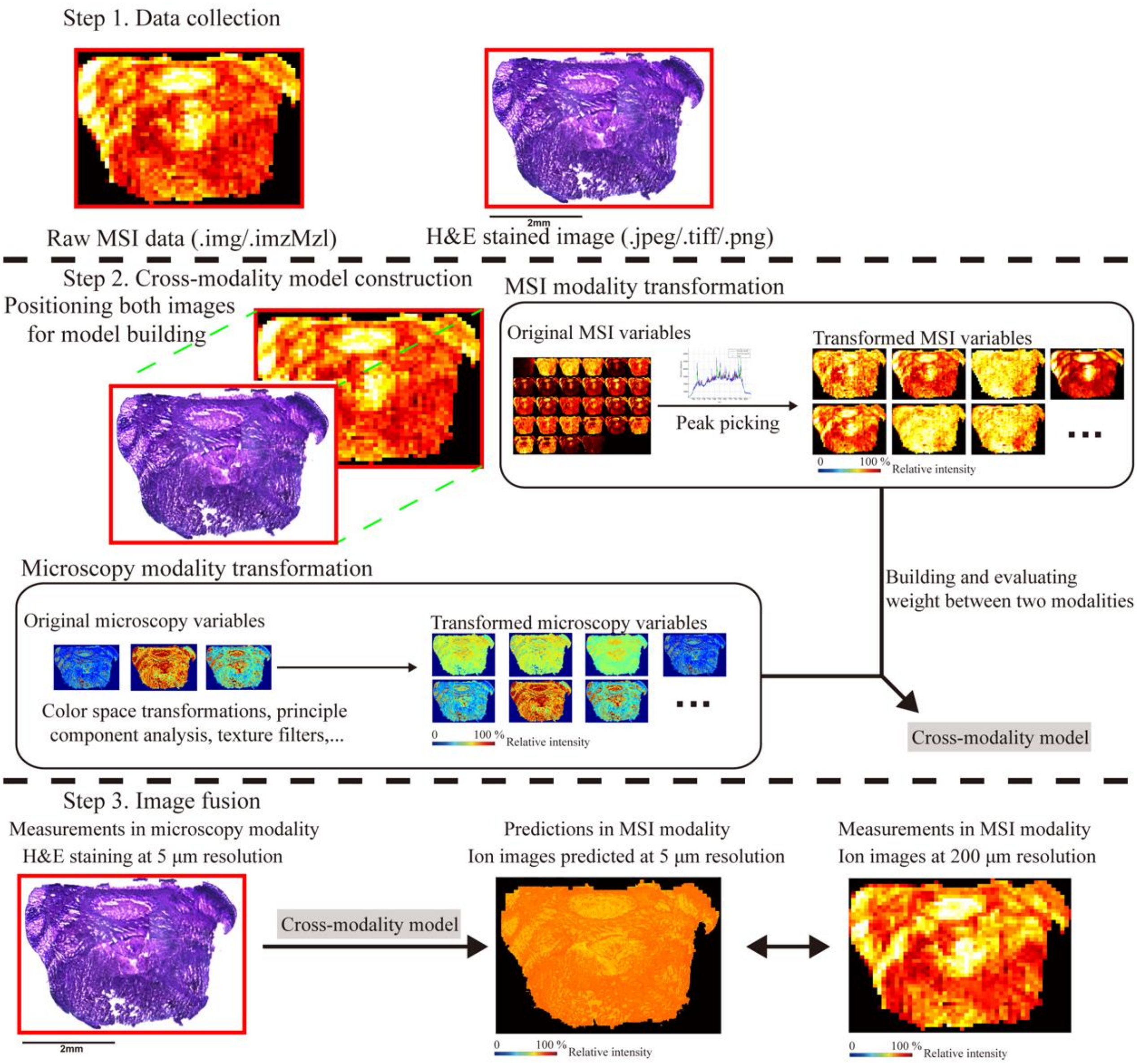


**Supplementary Figure 3.** Workflows and Protocols of ambient ionization MSI-microscopy image fusion. Raw mass spectra were collected by a Thermo LTQ Orbitrap hybrid mass spectrometer and converted into 2D raw MSI. Hematoxylin and eosin (H&E) staining was applied to scanned (for DESI) or adjacent tissue sections (for nanoDESI). Before image fusion, raw MSI and H&E stained photos were overlaid for position alignment. Detailed mass spectral information (including mass peaks, signal intensity, and position) and pixel RGB values of the images of H&E stained sections were then imported into the image fusion package for model building procedure. After model building, predicted high resolution MSI was be reconstructed.

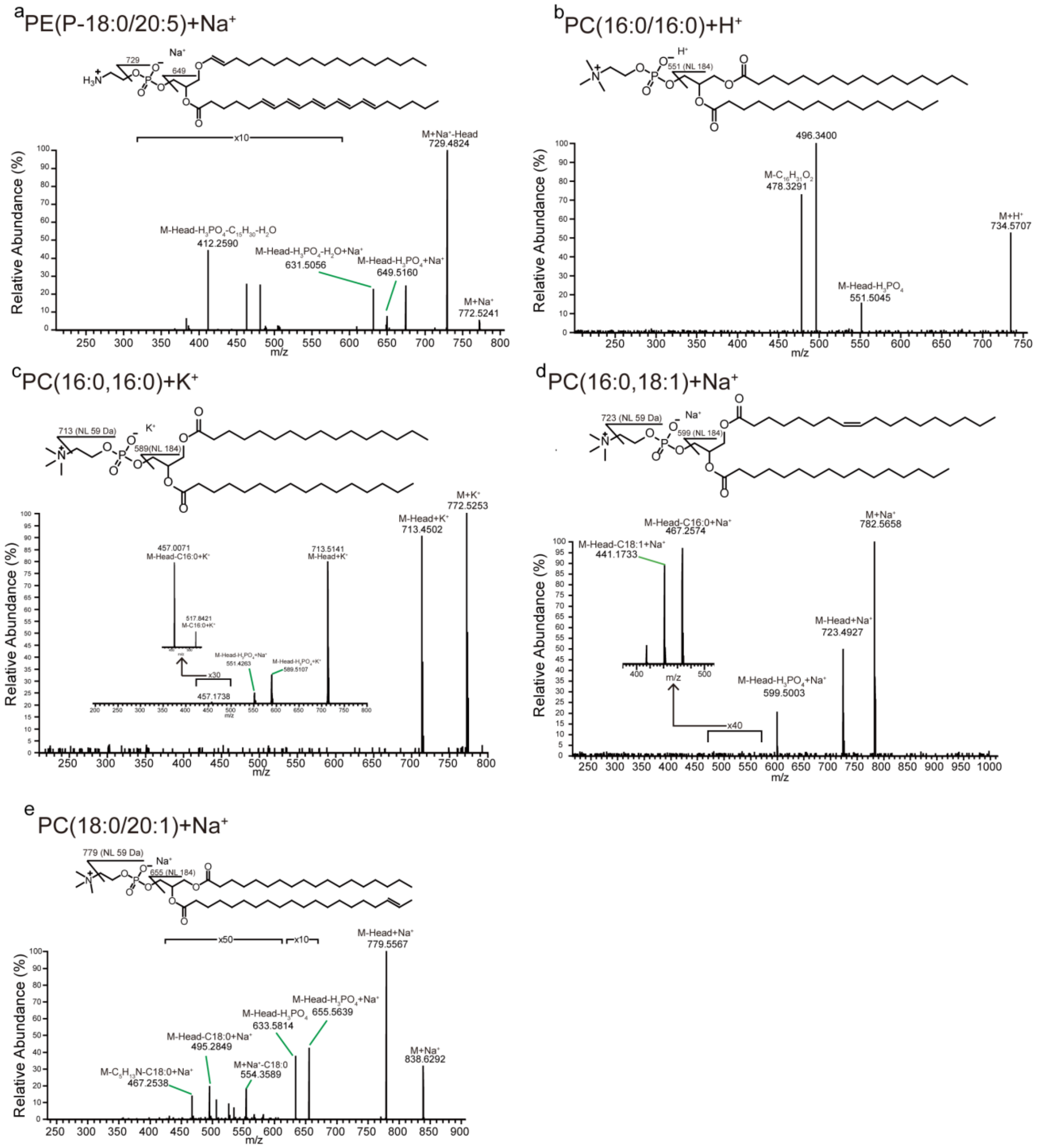


**Supplementary Figure 4.** Tandem mass analysis of annotated lipid species observed in mice tissue sections. (a) PE(P-18:0/20:5)+$Na^+$. (b) PC(16:0/16:0)+$H^+$. (b) PC(16:0/16:0)+$K^+$. (d) PC(16:0/18:1)+$Na^+$. (d) PC(18:0/20:1) +$Na^+$.

**Supplementary Table 1.** List of annotated phospholipids by tandem mass spectrometry analysis.

| Compound ID | Measured *m/z* | Theoretical *m/z* | Mass error (ppm) | Molecular formula | Adduct forms |
|---|---|---|---|---|---|
| PC(16:0/16:0) | 734.5707 | 734.5699 | -1.1 | $C_{40}H_{80}NO_8P$ | $[M+H]^+$ |
| PE(P-18/20:5) | 772.5246 | 772.5253 | -0.9 | $C_{43}H_{75}NO_7P$ | $[M+Na]^+$ |
| PC(18:1/16:0) | 782.5660 | 782.5670 | -1.3 | $C_{42}H_{82}NO_8P$ | $[M+Na]^+$ |
| PC(18:1/16:0) | 798.5407 | 798.5410 | -0.4 | $C_{42}H_{82}NO_8P$ | $[M+K]^+$ |
| PC(18:0/20:1) | 838.6292 | 838.6302 | -0.2 | $C_{46}H_{90}NO_8P$ | $[M+Na]^+$ |

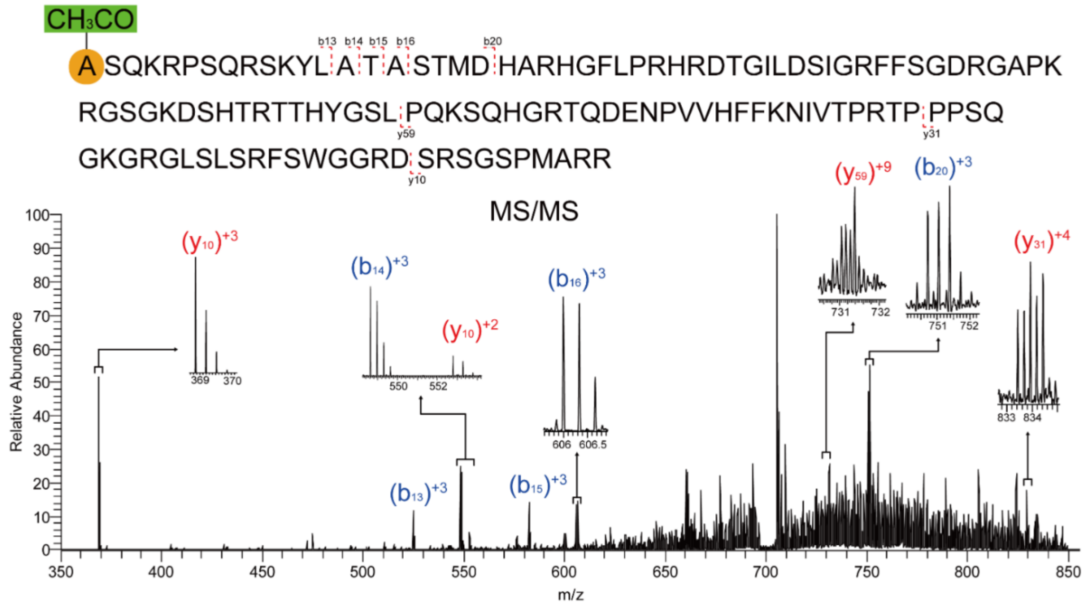


Number of Amino Acids: 127
Theoretical Mass: 14113.2 Da
Mass Difference: 0.184 Da
B Ions: 5 Y Ions: 3

| Name | Theoretical Mass (Da) | Observed Mass (Da) | Mass Difference (Da) | Mass Difference (ppm) |
|---|---|---|---|---|
| B13 | 1571.85315 | 1571.8503 | -0.00285 | -1.813146476 |
| B14 | 1642.89026 | 1642.8879 | -0.00236 | -1.436492782 |
| B15 | 1743.93794 | 1743.9342 | -0.00374 | -2.144571727 |
| B16 | 1814.97505 | 1814.9715 | -0.00355 | -1.955949752 |
| B20 | 2249.12219 | 2249.1192 | -0.00299 | -1.329407541 |
| Y10 | 1103.561805 | 1103.5587 | -0.003105 | -2.813616769 |
| Y31 | 3329.702395 | 3329.6856 | -0.016795 | -5.04399433 |
| Y59 | 6568.377165 | 6568.3773 | 0.000135 | 0.020553022 |

**Supplementary Figure 5.** Top-down tandem mass analysis of unmodified myelin basic protein (MBP) at *m/z* 707 (+20 charge).

MVLSGEDKSNIKAAWGKIGGHGAEYGAEALERMFASFPTTKTYFPHFDVSHGSAQVKGHG

KKVADALAXAAGHLDDLPGALSALSDLHAHKLRVDPVNFKLLSHCLLVTLASHHPAD

FTPAVHASLDKFLASVSTVLTSKYR

X = S(α1, 14944.7 Da)
T(α2, 14958.7 Da)
N(α3, 14971.7 Da)

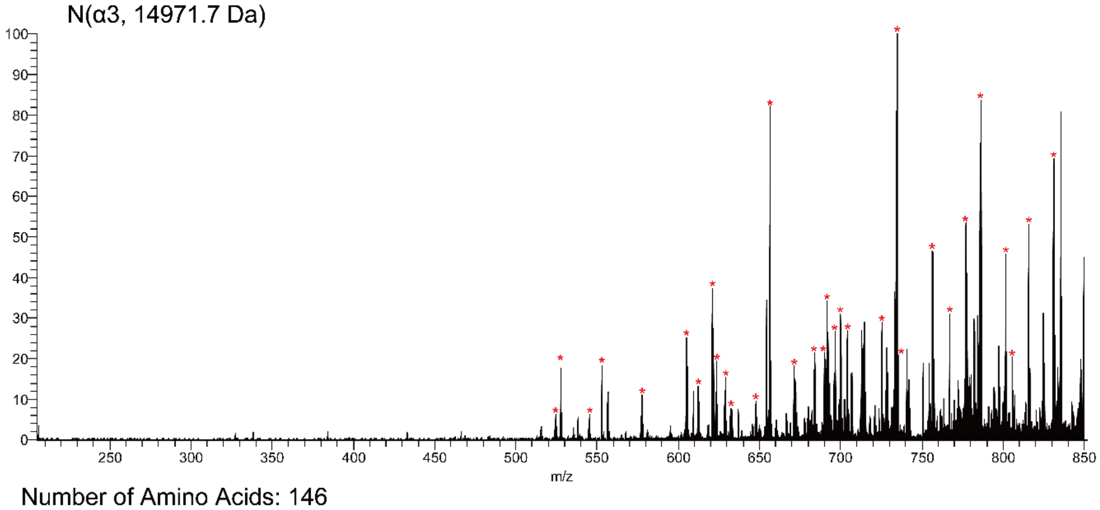


Number of Amino Acids: 146
Theoretical Mass: 14971.3 Da
Mass Difference: 2.44 Da
B Ions: 4 Y Ions: 32

| Name | Ion Type | Ion Number | Theoretical Mass | Observed Mass | Mass Difference (Da) | Mass Difference (ppm) |
|---|---|---|---|---|---|---|
| B22 | B | 22 | 2176.13884 | 2176.136 | -0.00284 | -1.305063789 |
| B23 | B | 23 | 2305.18143 | 2305.178 | -0.00343 | -1.487952295 |
| B75 | B | 75 | 7846.87661 | 7846.861 | -0.01561 | -1.989326553 |
| B77 | B | 77 | 8057.01343 | 8056.993 | -0.02043 | -2.53567903 |
| Y6 | Y | 6 | 766.433735 | 766.433 | -0.000735 | -0.958987015 |
| Y9 | Y | 9 | 1053.581855 | 1053.5804 | -0.001455 | -1.381003282 |
| Y10 | Y | 10 | 1152.650265 | 1152.6488 | -0.001465 | -1.270983962 |
| Y11 | Y | 11 | 1239.682295 | 1239.6808 | -0.001495 | -1.205954143 |
| Y12 | Y | 12 | 1310.719405 | 1310.7184 | -0.001005 | -0.766754499 |
| Y14 | Y | 14 | 1570.871875 | 1570.8702 | -0.001675 | -1.066286835 |
| Y18 | Y | 18 | 2014.109865 | 2014.1079 | -0.001965 | -0.975617087 |
| Y19 | Y | 19 | 2085.146975 | 2085.1449 | -0.002075 | -0.995133689 |
| Y20 | Y | 20 | 2222.205885 | 2222.2036 | -0.002285 | -1.02825756 |
| Y20 | Y | 20 | 2222.205885 | 2222.2044 | -0.001485 | -0.668254913 |
| Y23 | Y | 23 | 2489.364165 | 2489.3616 | -0.002565 | -1.030383596 |
| Y23 | Y | 23 | 2489.364165 | 2489.3619 | -0.002265 | -0.909870895 |
| Y28 | Y | 28 | 3020.597065 | 3020.5945 | -0.002565 | -0.849169865 |
| Y28 | Y | 28 | 3020.597065 | 3020.5956 | -0.001465 | -0.485003451 |
| Y29 | Y | 29 | 3157.655975 | 3157.652 | -0.003975 | -1.258845179 |
| Y32 | Y | 32 | 3452.784025 | 3452.7795 | -0.004525 | -1.310536648 |
| Y33 | Y | 33 | 3565.868085 | 3565.8645 | -0.003585 | -1.00536529 |
| Y34 | Y | 34 | 3666.915765 | 3666.9084 | -0.007365 | -2.008499915 |
| Y34 | Y | 34 | 3666.915765 | 3666.911 | -0.004765 | -1.299457175 |
| Y35 | Y | 35 | 3765.984175 | 3765.9795 | -0.004675 | -1.241375371 |
| Y35 | Y | 35 | 3765.984175 | 3765.9804 | -0.003775 | -1.002394016 |
| Y36 | Y | 36 | 3879.068235 | 3879.063 | -0.005235 | -1.349550893 |
| Y36 | Y | 36 | 3879.068235 | 3879.0655 | -0.002735 | -0.705066226 |
| Y38 | Y | 38 | 4095.161485 | 4095.1554 | -0.006085 | -1.48589989 |
| Y47 | Y | 47 | 5130.748015 | 5130.7417 | -0.006315 | -1.230814685 |
| Y56 | Y | 56 | 6200.362485 | 6200.3502 | -0.012285 | -1.981335774 |
| Y58 | Y | 58 | 6402.421455 | 6402.4136 | -0.007855 | -1.226879557 |
| Y59 | Y | 59 | 6515.505515 | 6515.4942 | -0.011315 | -1.736626571 |
| Y59 | Y | 59 | 6515.505515 | 6515.4992 | -0.006315 | -0.969226407 |
| Y60 | Y | 60 | 6586.542625 | 6586.5438 | 0.001175 | 0.178394048 |
| Y61 | Y | 61 | 6673.574655 | 6673.556 | -0.018655 | -2.79535346 |
| Y65 | Y | 65 | 7011.770045 | 7011.778 | 0.007955 | 1.134520948 |

**Supplementary Figure 6.** Top-down tandem mass analysis of Hemoglobin subunit alpha at *m/z* 751 (+ 20 charge).

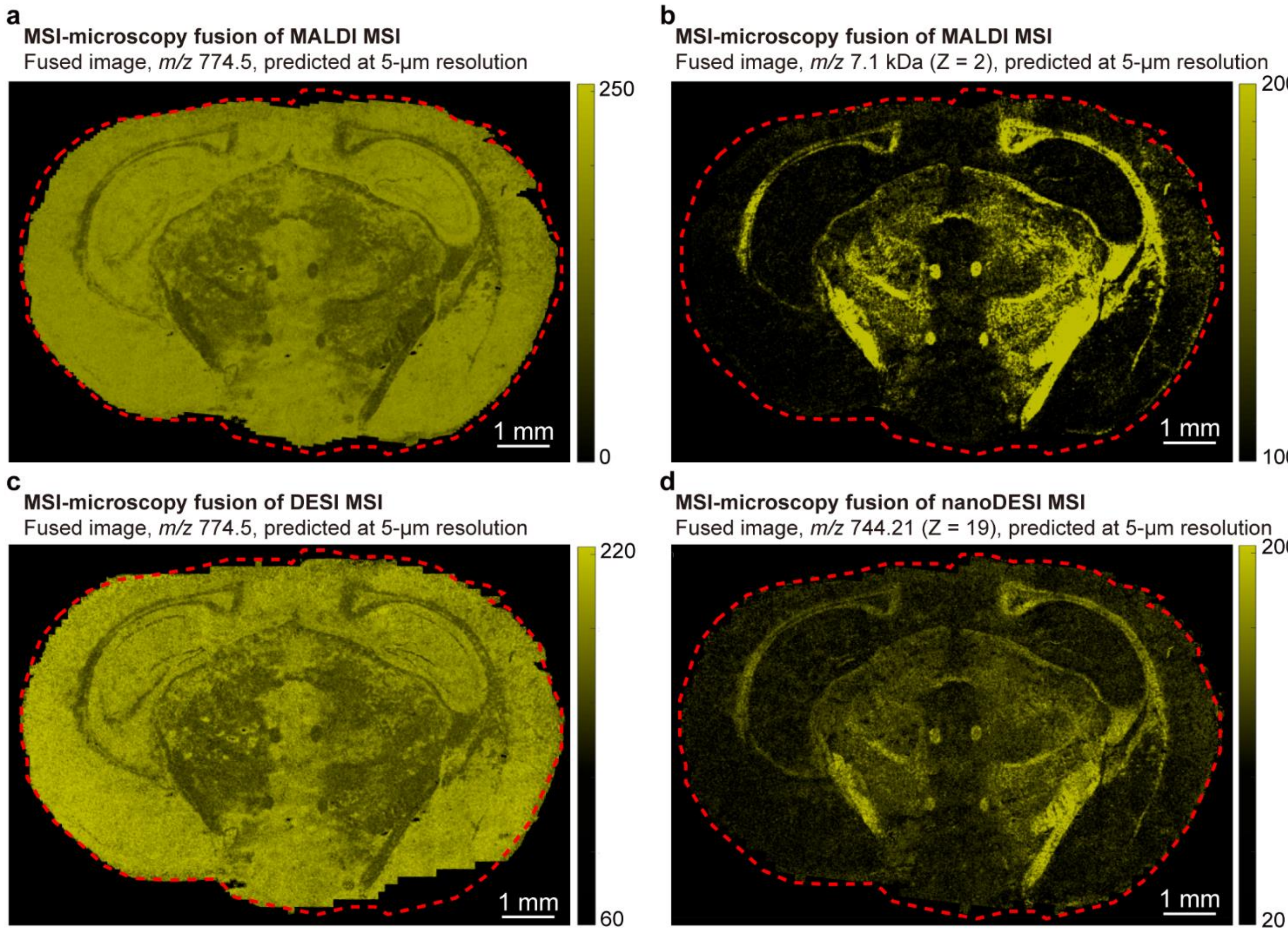


**Supplementary Figure 7.** Comparisons of image fusion of microscopy image with MALDI-TOF MSI and ambient ionization MSI. The MALDI-TOF MSI of the neighboring tissue sections (main text Fig. 4) targeting the same ions as DESI/nanoDESI MSI were also conducted with microscopy image fusion. (a) MALDI-TOF MSI-microscopy fusion image of lipids of *m/z* 774.5 (b) MALDI-TOF MSI-microscopy fusion image of protein of *m/z* 7060 (+2 charge). These results are highly similar to the results from the predictive results for (c) DESI and (d) nanoDESI MSI.

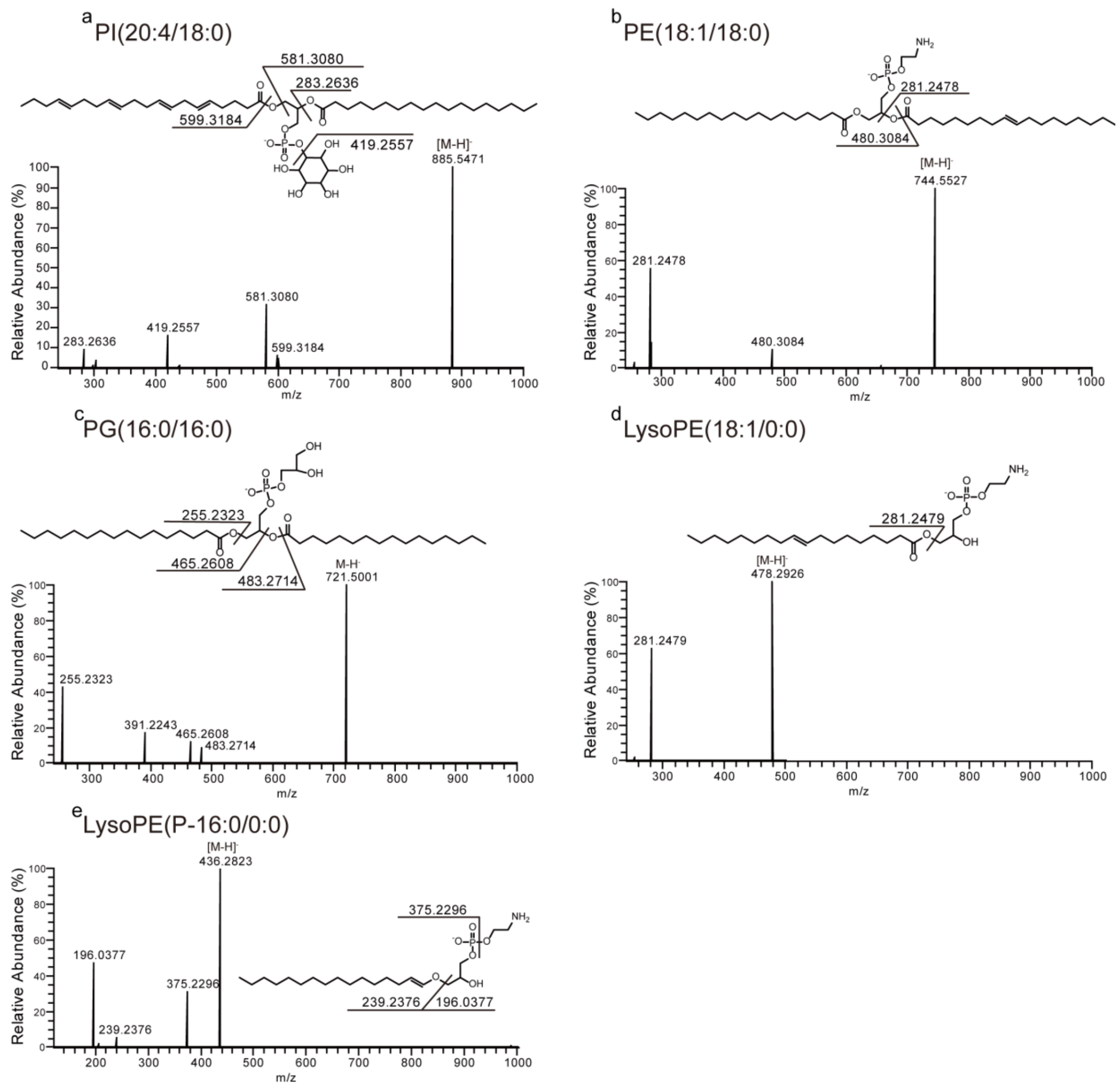


**Supplementary Figure 8.** Tandem mass analysis of lipids in the metastatic mouse lung tissue section, including (a) PI(20:4/18:0)-H$^-$. (b) PI(18:1/18:0)-H$^-$. (c) PG(16:0/16:0)-H$^-$. (d) LysoPE(18:1/0:0) -H$^-$. (d) LysoPE(P-16:0/0:0) -H$^-$.

**Supplementary Table 2.** List of annotations for representative potential lipid biomarkers for cancer diagnosis.

| Compound # | Measured *m/z* | Theoretical *m/z* | Mass error (ppm) | Molecular formula | Adduct form |
|---|---|---|---|---|---|
| PI(20:4/18:0) | 885.5471 | 885.5499 | -3.1 | $C_{47}H_{83}O_{13}P$ | $[M-H]^-$ |
| PE(18:1/18:0) | 744.5527 | 744.5548 | -2.8 | $C_{41}H_{80}NO_8P$ | $[M-H]^-$ |
| PE(18:1/0:0) | 478.2926 | 478.2939 | -2.7 | $C_{23}H_{46}NO_7P$ | $[M-H]^-$ |
| PE(P-16:0/0:0) | 436.2823 | 436.2833 | -2.2 | $C_{21}H_{44}NO_6P$ | $[M-H]^-$ |

### Lipids were annotated by tandem mass spectrometry analysis using DESI source for direct interrogations on the surface tissue section.

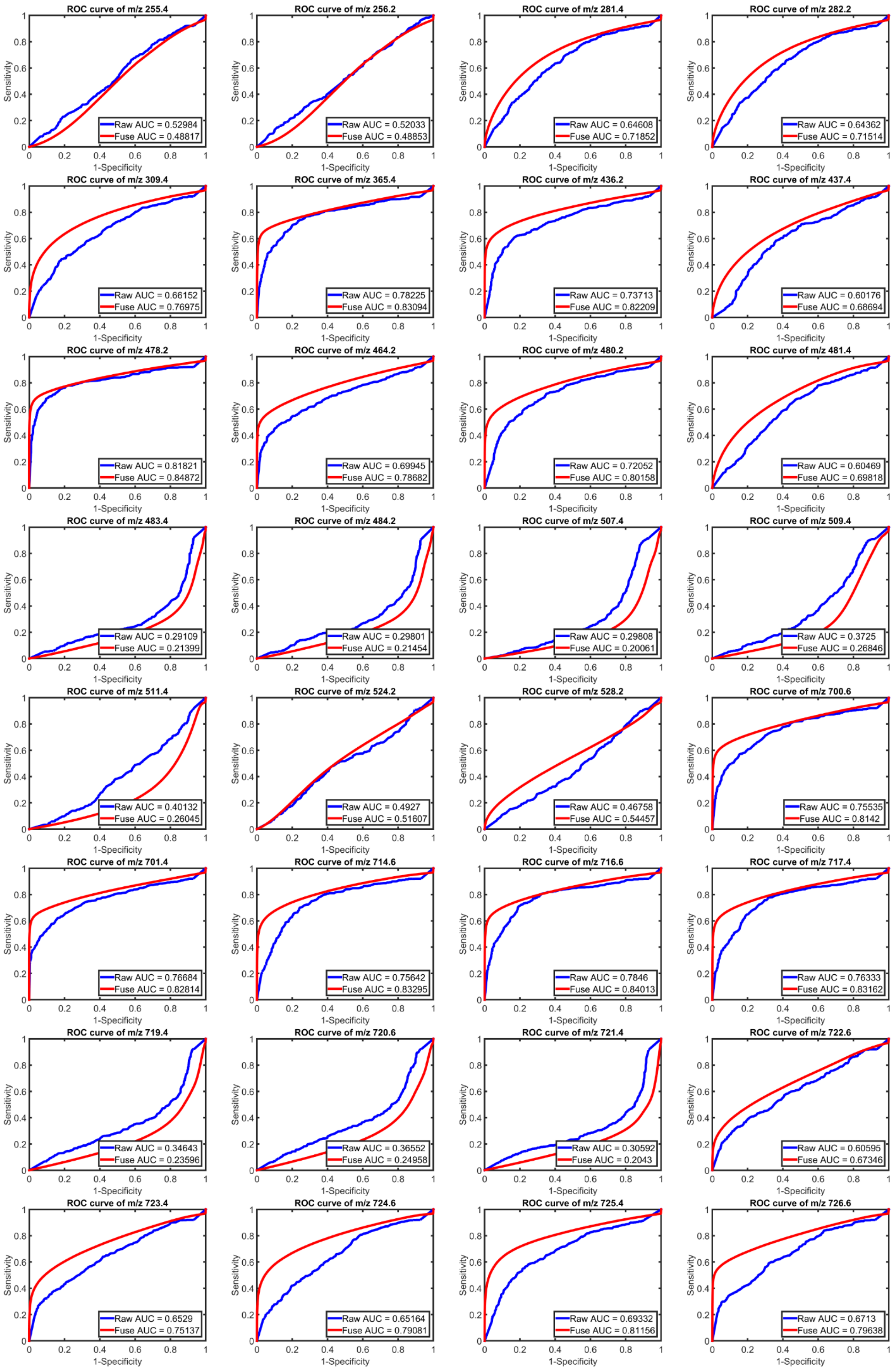
ROC curve of m/z 255.4
Raw AUC = 0.52984
Fuse AUC = 0.48817
ROC curve of m/z 256.2
Raw AUC = 0.52033
Fuse AUC = 0.48853
ROC curve of m/z 281.4
Raw AUC = 0.64608
Fuse AUC = 0.71852
ROC curve of m/z 282.2
Raw AUC = 0.64362
Fuse AUC = 0.71514
ROC curve of m/z 309.4
Raw AUC = 0.66152
Fuse AUC = 0.76975
ROC curve of m/z 365.4
Raw AUC = 0.78225
Fuse AUC = 0.83094
ROC curve of m/z 436.2
Raw AUC = 0.73713
Fuse AUC = 0.82209
ROC curve of m/z 437.4
Raw AUC = 0.60176
Fuse AUC = 0.68694
ROC curve of m/z 478.2
Raw AUC = 0.81821
Fuse AUC = 0.84872
ROC curve of m/z 464.2
Raw AUC = 0.69945
Fuse AUC = 0.78682
ROC curve of m/z 480.2
Raw AUC = 0.72052
Fuse AUC = 0.80158
ROC curve of m/z 481.4
Raw AUC = 0.60469
Fuse AUC = 0.69818
ROC curve of m/z 483.4
Raw AUC = 0.29109
Fuse AUC = 0.21399
ROC curve of m/z 484.2
Raw AUC = 0.29801
Fuse AUC = 0.21454
ROC curve of m/z 507.4
Raw AUC = 0.29808
Fuse AUC = 0.20061
ROC curve of m/z 509.4
Raw AUC = 0.3725
Fuse AUC = 0.26846
ROC curve of m/z 511.4
Raw AUC = 0.40132
Fuse AUC = 0.26045
ROC curve of m/z 524.2
Raw AUC = 0.4927
Fuse AUC = 0.51607
ROC curve of m/z 528.2
Raw AUC = 0.46758
Fuse AUC = 0.54457
ROC curve of m/z 700.6
Raw AUC = 0.75535
Fuse AUC = 0.8142
ROC curve of m/z 701.4
Raw AUC = 0.76684
Fuse AUC = 0.82814
ROC curve of m/z 714.6
Raw AUC = 0.75642
Fuse AUC = 0.83295
ROC curve of m/z 716.6
Raw AUC = 0.7846
Fuse AUC = 0.84013
ROC curve of m/z 717.4
Raw AUC = 0.76333
Fuse AUC = 0.83162
ROC curve of m/z 719.4
Raw AUC = 0.34643
Fuse AUC = 0.23596
ROC curve of m/z 720.6
Raw AUC = 0.36552
Fuse AUC = 0.24958
ROC curve of m/z 721.4
Raw AUC = 0.30592
Fuse AUC = 0.2043
ROC curve of m/z 722.6
Raw AUC = 0.60595
Fuse AUC = 0.67346
ROC curve of m/z 723.4
Raw AUC = 0.6529
Fuse AUC = 0.75137
ROC curve of m/z 724.6
Raw AUC = 0.65164
Fuse AUC = 0.79081
ROC curve of m/z 725.4
Raw AUC = 0.69332
Fuse AUC = 0.81156
ROC curve of m/z 726.6
Raw AUC = 0.6713
Fuse AUC = 0.79638
Sensitivity
1-Specificity

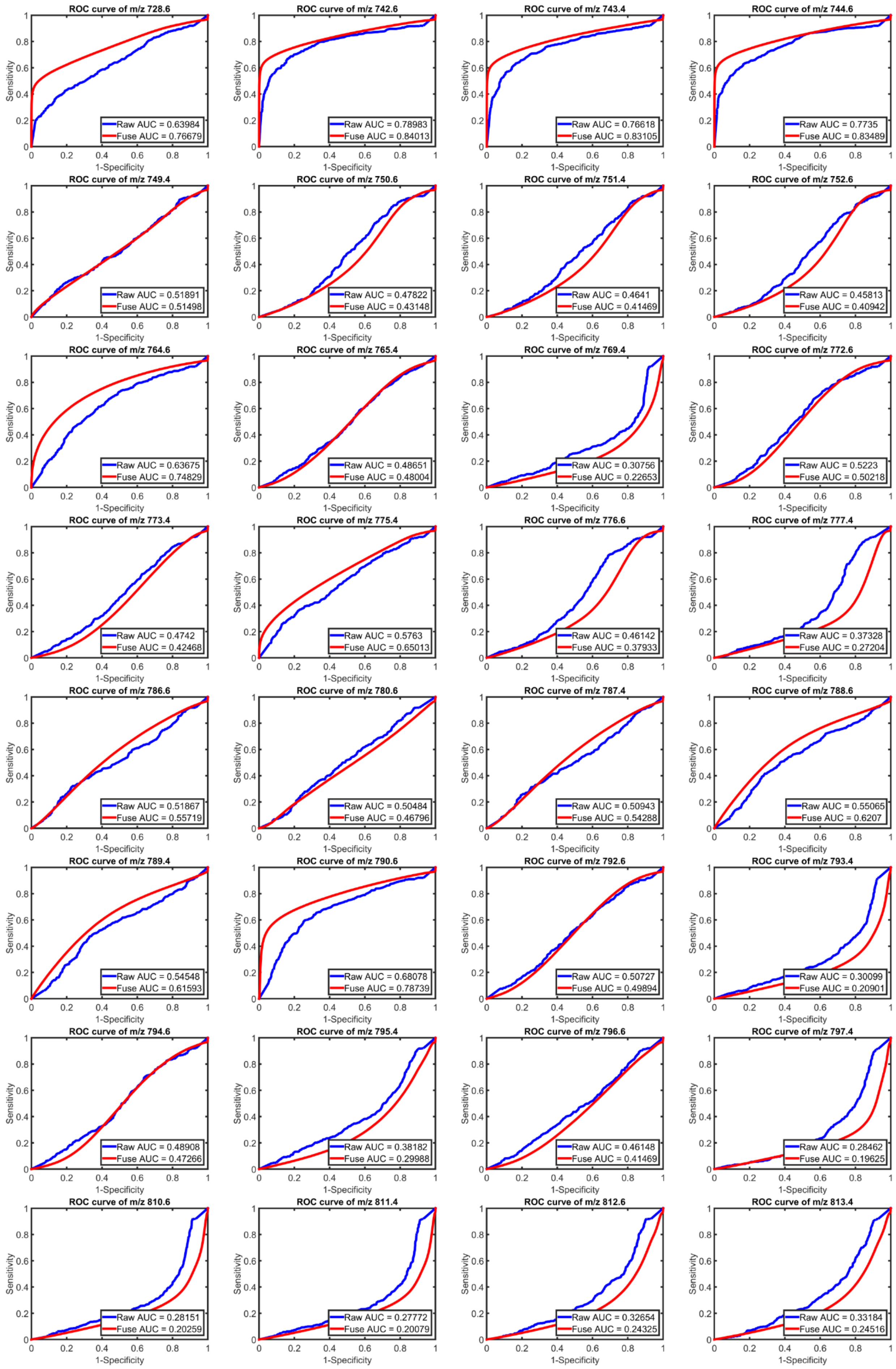
ROC curve of m/z 728.6
Raw AUC = 0.63984
Fuse AUC = 0.76679
ROC curve of m/z 742.6
Raw AUC = 0.78983
Fuse AUC = 0.84013
ROC curve of m/z 743.4
Raw AUC = 0.76618
Fuse AUC = 0.83105
ROC curve of m/z 744.6
Raw AUC = 0.7735
Fuse AUC = 0.83489
ROC curve of m/z 749.4
Raw AUC = 0.51891
Fuse AUC = 0.51498
ROC curve of m/z 750.6
Raw AUC = 0.47822
Fuse AUC = 0.43148
ROC curve of m/z 751.4
Raw AUC = 0.4641
Fuse AUC = 0.41469
ROC curve of m/z 752.6
Raw AUC = 0.45813
Fuse AUC = 0.40942
ROC curve of m/z 764.6
Raw AUC = 0.63675
Fuse AUC = 0.74829
ROC curve of m/z 765.4
Raw AUC = 0.48651
Fuse AUC = 0.48004
ROC curve of m/z 769.4
Raw AUC = 0.30756
Fuse AUC = 0.22653
ROC curve of m/z 772.6
Raw AUC = 0.5223
Fuse AUC = 0.50218
ROC curve of m/z 773.4
Raw AUC = 0.4742
Fuse AUC = 0.42468
ROC curve of m/z 775.4
Raw AUC = 0.5763
Fuse AUC = 0.65013
ROC curve of m/z 776.6
Raw AUC = 0.46142
Fuse AUC = 0.37933
ROC curve of m/z 777.4
Raw AUC = 0.37328
Fuse AUC = 0.27204
ROC curve of m/z 786.6
Raw AUC = 0.51867
Fuse AUC = 0.55719
ROC curve of m/z 780.6
Raw AUC = 0.50484
Fuse AUC = 0.46796
ROC curve of m/z 787.4
Raw AUC = 0.50943
Fuse AUC = 0.54288
ROC curve of m/z 788.6
Raw AUC = 0.55065
Fuse AUC = 0.6207
ROC curve of m/z 789.4
Raw AUC = 0.54548
Fuse AUC = 0.61593
ROC curve of m/z 790.6
Raw AUC = 0.68078
Fuse AUC = 0.78739
ROC curve of m/z 792.6
Raw AUC = 0.50727
Fuse AUC = 0.49894
ROC curve of m/z 793.4
Raw AUC = 0.30099
Fuse AUC = 0.20901
ROC curve of m/z 794.6
Raw AUC = 0.48908
Fuse AUC = 0.47266
ROC curve of m/z 795.4
Raw AUC = 0.38182
Fuse AUC = 0.29988
ROC curve of m/z 796.6
Raw AUC = 0.46148
Fuse AUC = 0.41469
ROC curve of m/z 797.4
Raw AUC = 0.28462
Fuse AUC = 0.19625
ROC curve of m/z 810.6
Raw AUC = 0.28151
Fuse AUC = 0.20259
ROC curve of m/z 811.4
Raw AUC = 0.27772
Fuse AUC = 0.20079
ROC curve of m/z 812.6
Raw AUC = 0.32654
Fuse AUC = 0.24325
ROC curve of m/z 813.4
Raw AUC = 0.33184
Fuse AUC = 0.24516
Sensitivity
1-Specificity

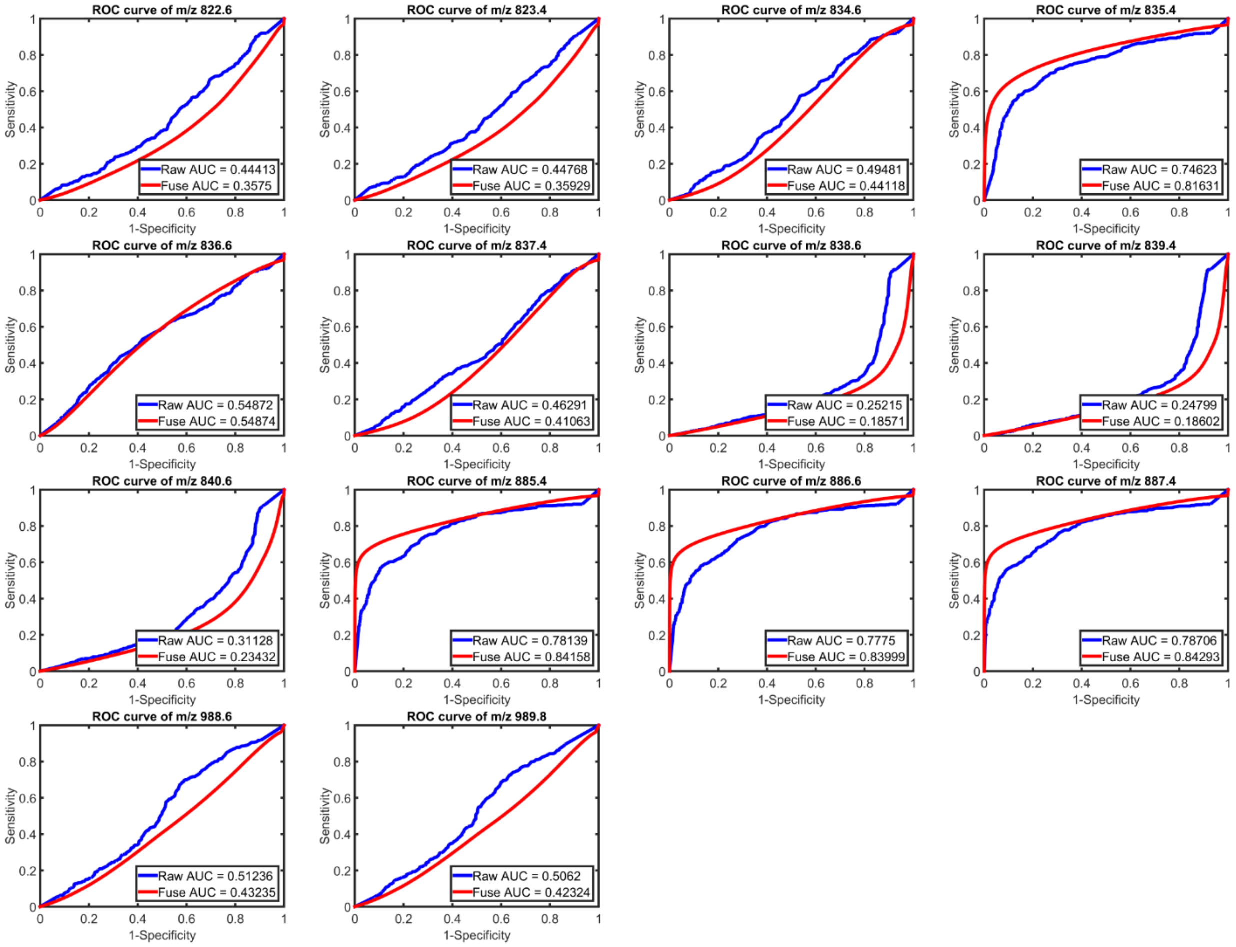


**Supplementary Figure 9.** ROC curves of individual ions with different *m/z* observed on the metastatic mouse lung tissue section and their corresponding AUC. The sensitivity (true positive rate) was defined as the ratio of "the number of Cancer pixels determined by both MSI and the pathologist" to "the number of Cancer pixels labeled by the pathologist"; the (1-specificity), or false positive rate, was defined as the ratio of "the number of Cancer pixels determined by MSI in the normal tissue region" to "the number Normal pixels labeled by the pathologist". The ROC curves of an *m/z* peak were sketched by setting the intensity threshold from 0% to 100% of the highest intensity of the ion in all pixels.

**Supplementary Table 3.** List of potential biomarkers for mouse lung cancer diagnosis before & after image fusion[#].

| **Potential biomarkers from raw DESI MSI** | | **Potential biomarkers from predicted MSI** | |
|---|---|---|---|
| ***m/z* *** | **Area under curve (AUC)** | ***m/z* *** | **Area under curve (AUC)** |
| | | 281.4 | 0.72 |
| | | 282.2 | 0.72 |
| | | 309.4 | 0.77 |
| 365.4 | 0.78 | 365.4 | 0.83 |
| 436.2 | 0.74 | 436.2 | 0.82 |
| | | 464.2 | 0.79 |
| 478.2 | 0.82 | 478.2 | 0.85 |
| 480.2 | 0.72 | 480.2 | 0.80 |
| 700.6 | 0.76 | 700.6 | 0.81 |
| 701.4 | 0.77 | 701.4 | 0.83 |
| 714.6 | 0.76 | 714.6 | 0.83 |
| 716.6 | 0.78 | 716.6 | 0.84 |
| 717.4 | 0.76 | 717.4 | 0.83 |
| | | 723.4 | 0.75 |
| | | 724.6 | 0.79 |
| | | 725.4 | 0.81 |
| | | 726.6 | 0.80 |
| | | 728.6 | 0.77 |
| 742.6 | 0.79 | 742.6 | 0.84 |
| 743.4 | 0.77 | 743.4 | 0.83 |
| 744.6 | 0.77 | 744.6 | 0.83 |
| | | 764.6 | 0.75 |
| | | 790.6 | 0.79 |
| 835.4 | 0.75 | 835.4 | 0.82 |
| 885.4 | 0.78 | 885.4 | 0.84 |
| 886.6 | 0.78 | 886.6 | 0.84 |
| 887.4 | 0.79 | 887.4 | 0.84 |

[#] The potential biomarkers are defined as the ions with AUC of the ROC curves > 0.7.

* The *m/z* values were binned every 0.4 Da.

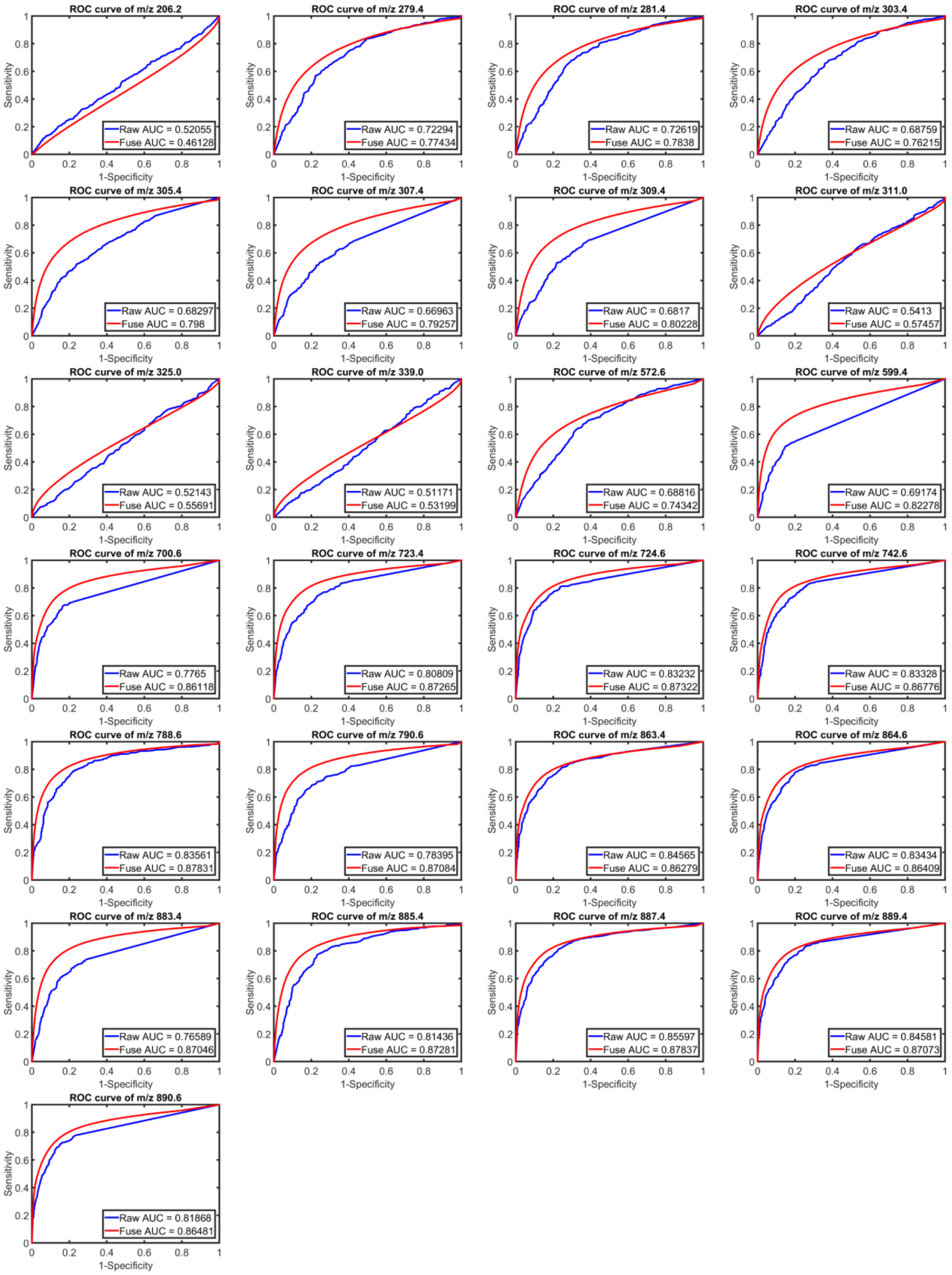


**Supplementary Figure 10.** ROC curves of individual ions with different *m/z* observed on the Luminal B human breast cancer tissue section and their corresponding AUC. The sensitivity (true positive rate) was defined as the ratio of “the number of Cancer pixels determined by both MSI and the pathologist” to “the number of Cancer pixels labeled by the pathologist”; the (1-specificity), or false positive rate, was defined as the ratio of “the number of Cancer pixels determined by MSI in the normal tissue region” to “the number Normal pixels labeled by the pathologist”. The ROC curves of an *m/z* peak were sketched by setting the intensity threshold from 0% to 100% of the highest intensity of the ion in all pixels.

**Supplementary Table 4.** List of potential biomarkers for human breast cancer diagnosis before & after image fusion[#].

| Potential biomarkers from raw DESI MSI | | Potential biomarkers from predicted MSI | |
|---|---|---|---|
| ***m/z* *** | **Area under curve (AUC)** | ***m/z* *** | **Area under curve (AUC)** |
| 279.4 | 0.72 | 279.4 | 0.77 |
| 281.4 | 0.73 | 281.4 | 0.78 |
| | | 303.4 | 0.76 |
| | | 305.4 | 0.80 |
| | | 307.4 | 0.79 |
| | | 309.4 | 0.80 |
| | | 572.6 | 0.74 |
| | | 599.4 | 0.82 |
| 700.6 | 0.78 | 700.6 | 0.86 |
| 723.4 | 0.81 | 723.4 | 0.87 |
| 724.6 | 0.83 | 724.6 | 0.87 |
| 742.6 | 0.83 | 742.6 | 0.87 |
| 747.4 | 0.73 | 747.4 | 0.86 |
| 760.6 | 0.82 | 760.6 | 0.87 |
| 766.6 | 0.81 | 766.6 | 0.87 |
| 786.6 | 0.84 | 786.6 | 0.87 |
| 788.6 | 0.84 | 788.6 | 0.88 |
| 790.6 | 0.78 | 790.6 | 0.87 |
| 809.4 | 0.79 | 809.4 | 0.85 |
| 810.6 | 0.82 | 810.6 | 0.88 |
| 812.6 | 0.83 | 812.6 | 0.88 |
| 835.4 | 0.84 | 835.4 | 0.86 |
| 837.4 | 0.82 | 837.4 | 0.86 |
| 838.6 | 0.83 | 838.6 | 0.87 |
| 859.4 | 0.79 | 859.4 | 0.87 |
| 861.4 | 0.85 | 861.4 | 0.87 |
| 863.4 | 0.85 | 863.4 | 0.86 |
| 864.6 | 0.83 | 864.6 | 0.86 |
| 883.4 | 0.77 | 883.4 | 0.87 |
| 885.4 | 0.81 | 885.4 | 0.87 |
| 887.4 | 0.86 | 887.4 | 0.88 |
| 889.4 | 0.85 | 889.4 | 0.87 |
| 890.6 | 0.82 | 890.6 | 0.86 |

[#] The potential biomarkers are defined as the ions with AUC of the ROC curves > 0.7.

* The *m/z* values were binned every 0.4 Da.
